# A transcriptional response to replication stress selectively expands a subset of *BRCA2*-mutant mammary epithelial cells

**DOI:** 10.1101/2022.11.14.516328

**Authors:** Maryam Ghaderi Najafabadi, G. Kenneth Gray, Li Ren Kong, Komal Gupta, David Perera, Joan Brugge, Ashok Venkitaraman, Mona Shehata

## Abstract

BRCA2 mutation carriers preferentially develop luminal-like breast cancers, but it remains unclear how BRCA2 mutations affect mammary epithelial subpopulations. Here, we report that Brca2^mut/WT^ mammary organoids subjected to replication stress activated a transcriptional response that selectively expands Brca2^mut/WT^ luminal cells lacking hormone receptor expression (HR-). While CyTOF analyses revealed comparable epithelial compositions among wildtype and Brca2^mut/WT^ mammary glands, Brca2^mut/WT^ HR- luminal cells exhibited greater organoid formation and preferentially survived and expanded under replication stress. ScRNA-seq analysis corroborated the expansion of HR- luminal cells which express elevated levels of Tetraspanin-8 (*Tspan8*) and *Thrsp* mRNA, and pathways implicated in replication stress survival including Type I interferon responses. Notably, CRISPR/Cas9-mediated deletion of *Tspan8* or *Thrsp* prevented Brca2^mut/WT^ HR- luminal cell expansion. Our findings indicate that Brca2^mut/WT^ cells have an activate a transcriptional response after replication stress that preferentially favours outgrowth of HR- luminal cells through the expression of interferon-responsive and mammary alveolar genes.

## Introduction

Women who inherit pathogenic germline BRCA2 mutations exhibit increased risk of developing luminal subtype breast cancers^1,2^. What drives the development of these malignancies remains unclear. One possibility is that germline BRCA2 mutations cause defects in the differentiation or outgrowth of selected mammary epithelial subpopulations. This possibility has been best studied in germline BRCA1-assocatied tumours, a related tumour suppressor gene whose inactivation predisposes women to triple negative breast cancer subtypes. BRCA1 mutant breast tumours originate from specific cells-of-origin^3,4^, and germline BRCA1 carriers contain expanded mammary luminal progenitor populations compared to non-carrier individuals prior to cancer onset^3,5^.

However, there is relatively little information concerning the steps that promote breast cancer development in *BRCA2* mutation carriers, and even less information regarding the BRCA2 mutational effects on the different mammary epithelial subpopulations. For example, myoepithelial cell populations in samples from BRCA2 carriers showed no phenotype alterations^6^. For luminal cell populations, conflicting evidence exists regarding progenitor capacity in BRCA2 carriers^5,7,8^. Throughout the reproductive stages, murine models bearing Brca2 monoallelic germline mutations^9^ or hypomorphic biallelic Brca2 mutations^10^ display normal mammary glands, suggesting that mammary lineage trajectory is unperturbed by *Brca2* heterozygosity or haploinsufficiency.

There is compelling evidence that BRCA2 is an essential component of the cellular response to DNA replication stress. Cells lacking Brca2 exhibit chromosomal instability during cell division^11,12^ accompanied by defective DNA repair by homologous DNA recombination^13^. Mammary myoepithelial and luminal cells lacking Brca2 exhibit defects in DNA repair by homologous recombination^10^. Cells bearing monoallelic Brca2 mutations retain these functions^10–12^. How mammary cells in BRCA2 mutation carriers respond to replication stress is unclear. As all mammary epithelial lineages have proliferative ability^14–16^, and therefore experience replication stress. Yet, clinical evidence demonstrates that BRCA2 carriers tend to develop luminal-like breast cancers^1,2^. These observations raise the possibility that monoallelic Brca2 mutations may differentially affect replication stress responses in different mammary epithelial subtypes.

In this study, we investigated the effect of replication stress on mammary organoids derived from transgenic strains bearing a monoallelic truncating *Brca2* mutation (Brca2^mut/WT^) compared with wildtype controls (wildtype). Histology and CyTOF analyses showed a similar proportional representation of epithelial subtypes in wildtype and Brca2^mut/WT^ mammary glands. Mammary organoid assays revealed organoid formation capacity was increased in Brca2^mut/WT^ HR- luminal cell subpopulations. Single exposure of hydroxyurea (HU), to induce replication stress, showed wildtype and Brca2^mut/WT^ mammary cells responded in equivalent manners. However, prolonged HU exposure caused a selective expansion of the Brca2^mut/WT^ HR- luminal compartment. Single cell RNA sequencing (scRNA-seq) analyses revealed upregulation of interferon response genes and an increase in the HR- luminal cycling cell cluster proportion. This cell cluster contained transcripts of *Tspan8* and *Thrsp*, both of which are expressed in mammary luminal cells^17,18^. Exposure to inhibitors, or deletion of the HR- luminal cycling cluster via CRISPR/Cas9-mediated Tspan8 and Thrsp knockout demonstrated that Brca2 heterozygous cells activate these pathways to circumvent replication-stress induced cell death and promote cell proliferative survival.

## Results

### Brca2^mut/WT^ mammary HR- luminal cells have increased organoid formation capacity

BRCA2 carriers develop functional breast tissues containing all epithelial subpopulations^5,7,8^, although changes in epithelial dynamics have been observed in some BRCA2 carriers^7^. Many pathogenic germline mutations in BRCA2 have been reported [Clinvar database (https://www.ncbi.nlm.nih.gov/clinvar/)], each with potentially heterogenous effects on biological function. Prior studies on human breast samples from mutation carriers have typically involved small numbers of comparable genotypes, however clinical samples have been derived from patients at different ages with distinct reproductive and clinical histories. For these reasons, it has been difficult to determine whether observed changes in epithelial dynamics are due to BRCA2 mutation, or other factors. To address these issues, we utilised a genetically engineered murine model carrying a monoallelic germline Brca2 truncating mutation, Brca2^mut/WT19^, which recapitulates pathogenic human variants in the so-called breast and ovarian cluster region of BRCA2. Brca2^mut/WT^ mice are previously reported to develop phenotypically normal mammary glands, based on gross examination^9,19^. We collected mammary glands from 3 young and 4 aged Brca2^mut/WT^ or wildtype mice. Histology sections show similar morphological appearance (Fig 1a), and the presence of myoepithelial and luminal lineages (Fig 1b, Supp Fig 1a).

**Figure 1:**
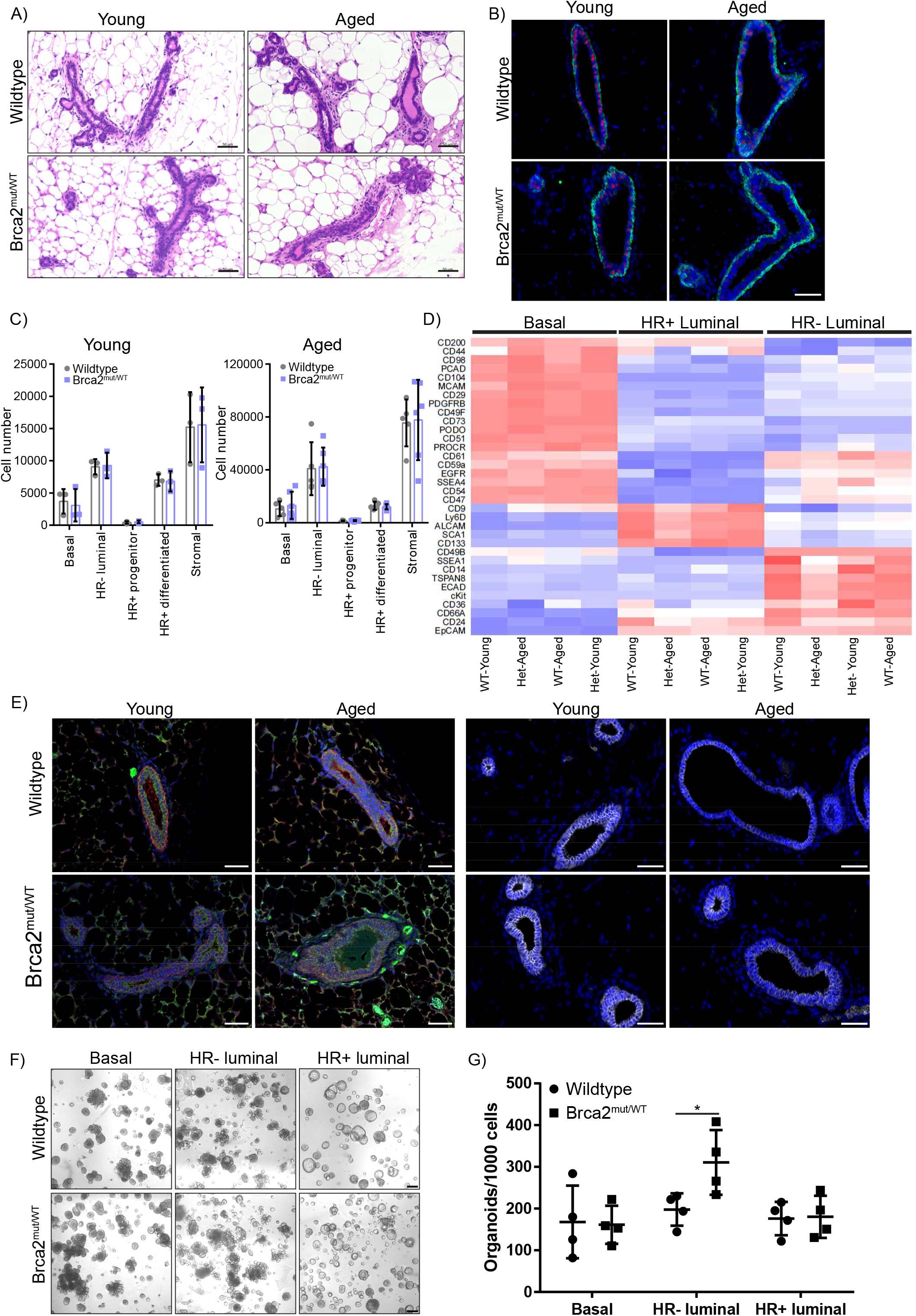
Brca2^mut/WT^ mammary glands are morphologically similar to wildtype mammary glands in young and aged mice. A) H&E staining of paraffin-embedded sections from young adult (3 months) or aged (8+ months) wildtype or Brca2^mut/WT^ mammary tissues. Images are representative of 3-4 mice/group. Scale bars, 50 µm. B) PR (magenta) and K14 (green) in young and aged wildtype or Brca2^mut/WT^ mammary glands. Images representative of 3-4 mice/group. Tissues were counter labelled with 4′,6-diamidino-2-phenylindole (DAPI, blue). Scale bars, 50 µm C) Absolute cell number of different mammary epithelial cell subpopulations of young or aged wildtype or Brca2^mut/WT^ mice as determined by flow cytometry. Mean +/- SD of 3-6 independent mice per group. An ANOVA followed by Tukey’s multiple comparison test showed no significant differences. D) Heat map showing expression levels of the indicated markers (y-axis) in the epithelial subpopulations from young and aged wildtype (WT) or Brca2^mut/WT^ (Het) mammary glands (red = high expression, blue = low expression). Data depicts the mean expression of 3 independent mice per group. E) Immunostaining of CD14 (green) and CD36 (red) (left) and E-Cadherin (grey, right) and DAPI (Blue) in young and aged wildtype and Brca2^mut/WT^ mammary glands. Images representative of 3-4 individual mice/group. Scale bars, 50 µm. F) Representative brightfield images of wildtype or Brca2^mut/WT^ organoids derived from Basal, HR- or HR+ luminal cells. Organoids derived from n=4 independent mice. Scale bar, 200 µm. G) Quantification of the number of wildtype or Brca2^mut/WT^ organoids formed from the different epithelial populations in F). Data presented as the mean +/- SD (n=4). Mann-Whitney 2 tailed unpaired test was performed, *p = 0.04.

To examine the epithelial proportions, we dissociated mammary glands from 3 young and 5-6 aged wildtype or Brca2^mut/WT^ mice and analysed the differential expression of CD49f and EpCAM using flow cytometry. The basal and stromal subpopulations contained comparable cell numbers between both genotypes in all ages (Fig 1c). The luminal population was further fractionated using expression of CD49b and Sca1 to distinguish HR+ and HR- luminal cells. No observable cell proportion differences were detected (Fig 1c). As the mammary gland is highly heterogenous with many complex cell states, we generated a mammary CyTOF mass cytometry panel consisting of 30-metal tagged antibodies, which recognise proteins associated with known mammary cell functions. CyTOF was carried out on oestrous matched young or aged wildtype and Brca2^mut/WT^ mammary glands.

Lineage relevant markers (Cd31, Cd45 and Ter119) were used to exclude haemopoietic, endothelial and stromal cells from the analysis. Based on the expression levels of the 30 markers, three clusters were evident, representing the three main epithelial lineages (Fig 1d). Basal and HR+ luminal subpopulations displayed no significant differences in protein expression irrespective of genotype or age. Several HR- luminal markers including CD61, SSEA-4, E-Cadherin and CD14 displayed slightly increased expression, in the Brca2^mut/WT^ aged HR- luminal populations compared to wildtype counterparts (Fig 1d); however, these were not statistically significant. Protein expression of CD14 and E-Cadherin on tissue sections further confirmed these results (Fig 1e, Supp Fig 1b-c). Thus, our findings indicate that loss of a single Brca2 allele in an unchallenged mammary gland does not affect the generation or maintenance of epithelial lineages.

Three dimensional (3D) organoids cultured from mammary tissues display a close resemblance to the epithelial composition of mammary glands, displaying budding structures that mimic branching morphogenesis^20^. A recently developed culture medium for mouse mammary organoid propagation has allowed organoids to be generated from a single cell. By seeding mammary single cell suspensions into an ECM (Matrigel) matrix (Supp Fig 1d), we established a diverse range of organoids, composed of spherical-like and multi-lobed structures (Supp Fig1e). Organoids started to form by day 5/6 and developed within 7-12 days (Supp fig1e). Wholemount staining confirmed the existence of multilineage organoids composed of a myoepithelial/basal outer layer (K14 expression) and an inner luminal layer containing both Progesterone receptor (PR) positive and negative luminal cells and are E-Cadherin positive (Supp Fig1f). Collectively, these findings demonstrate that a single mammary cell derived organoid recapitulates in-vivo mammary gland features.

Mammary glands from an aged cohort are predicted to be equivalent to a mid-aged human female^21^. Because the incidence of breast cancer in female BRCA2 mutation carriers increases at perimenopausal age, we used mammary glands from the aged cohorts for the remainder of this study to more closely model perimenopausal breast tissues. Only minimal changes in epithelial subtype proportions were observed, and so proliferation capacity of these subtypes was not examined. Next, we tested whether Brca2^mut/WT^ mammary epithelial cells have differential proliferative capacity. Glands from 4 aged mice (8-10 months) and basal, HR- and HR+ luminal populations were flow sorted and seeded into 3D cultures. All three populations were able to form 3D structures (Supp Fig1g). The basal and HR- luminal populations generated both spheroid-like and multi-lobed structures, indicating strong organoid formation ability (Fig 1f, Supp Fig 1g).

Wholemount staining confirmed the presence of both single lineage and multilineage organoids (Supp Fig 1h). The HR+ luminal population generated mainly spheroid-like structures, where the majority of organoids were PR+ indicating a lineage restricted cell type (Supp Fig 1h). While basal and HR+ luminal cells generated similar numbers of organoids from both genotypes (Fig 1g), Brca2^mut/WT^ HR- luminal cells displayed significantly increased organoid formation capacity (Fig 1g). In summary, we show that Brca2^mut/WT^ HR- luminal cells, while relatively equal in cell proportion, have greater proliferative capacity than their wildtype counterparts when propagated from single cells into organoids.

### Effect of short term replicative stress on wildtype and Brca2^mut/WT^ mammary organoids

To assess the effects of replication stress on the epithelial subpopulations from both genotypes, we established organoids from single mammary cells derived from 5-6 mice. On day 7, organoids were exposed to 1mM (Fig 2) or 4mM (Supp fig2) HU, which stalls DNA replication and assessed for γH2AX as a reporter of double strand DNA breaks, a consequence of replication stress (Fig 2a). As expected, γH2AX positive cells, measured by flow cytometry, was increased in the HU treated organoids. However, no differences were observed in γH2AX+ cells detected between the genotypes (Fig 2b, Supp Fig 2a), nor any changes in the proportion of the epithelial cell types after HU exposure (Fig 2c, Supp Fig 2b). These observations suggest that mammary epithelial cells carrying monoallelic Brca2 mutations retain a normal capacity to respond acutely to replication stress.

**Figure 2:**
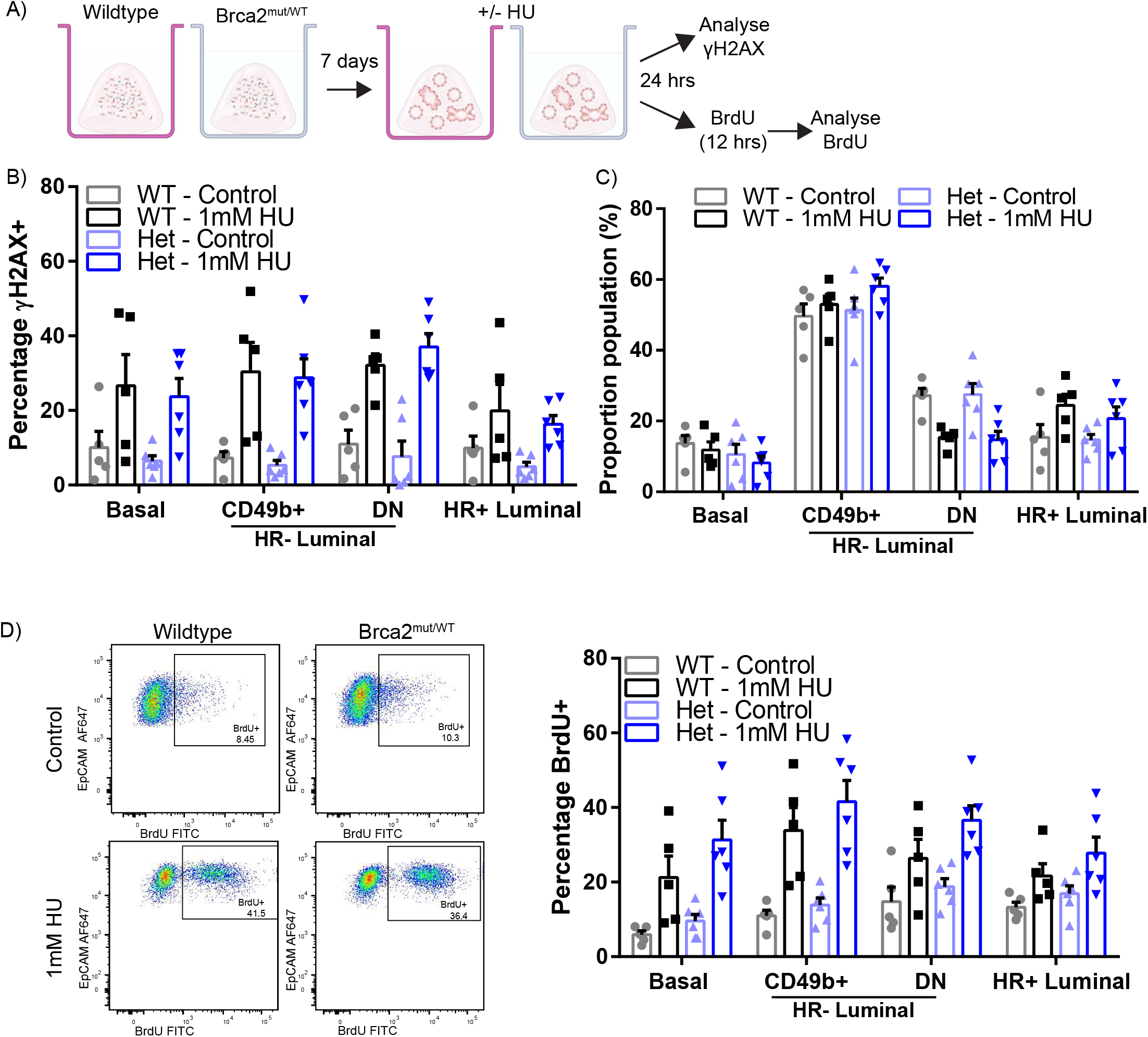
Brca2^mut/WT^ mammary organoids have similar response to short term DNA genotoxic stress as wildtype mammary organoids. A) Schematic illustrating the experimental pipeline for wildtype or Brca2^mut/WT^ cells grown as organoids, treated with 1mM HU and assessed for γH2AX or BrdU. B) Quantification of the percentage of γH2AX+ cells from wildtype or Brca2^mut/WT^ organoids post HU treatment. Data depicts the different epithelial populations. Data presented as the mean +/- SEM (n=5-6). C) Quantification of the proportion of epithelial populations in the wildtype or Brca2^mut/WT^ organoids post HU treatment. Data presented as mean +/- SEM (n=5-6). D) Representative flow cytometry analysis of BrdU+ CD49b+ luminal cells (left) and quantification (right) of the percentage of BrdU+ cells in the different epithelial subpopulations from wildtype or Brca2^mut/WT^ organoids post HU treatment. Data presented as mean +/- SEM (n=5-6). (B-D) An ANOVA followed by Fisher’s LSD test showed no significant differences.

To examine the effects of replication stress on cell proliferation, we administered BrdU to label the DNA of cells in the S phase of the cell cycle and assessed the proliferative activity across various cell populations (Fig 2a). Although the proportion of BrdU-labelled cells was increased in HU challenged organoids compared with unchallenged (Fig 2d), no significant differences in BrdU-labelled cells was detected between genotypes or epithelial lineages (Figure 2d, Supp Fig 2c). Thus, transient replicative stress evokes similar cellular phenotypes, irrespective of genotype in the short term.

### Expansion of a Brca2^mut/WT^ HR- luminal population after prolonged replication stress

However, replication stress in the breast may be prolonged rather than transient because the breast undergoes many proliferative and remodelling events over its lifetime. We therefore investigated the effects of prolonged exposures to replication stress in Brca2^mut/WT^ mammary organoids. Briefly, organoids (established from 5 mice per group) were exposed to 1mM HU for 24 hr and allowed to recover for 4 days, before passaging (Fig 3a). Organoids were exposed to HU in this way for 5 serial passages over 60 days (Fig 3a). Wildtype and mutant organoids not exposed to HU were used as controls. Staining for γH2AX after HU exposure confirmed that DNA damage was present at similar levels in all epithelial subpopulations over time (Supp Fig 3a).

**Figure 3:**
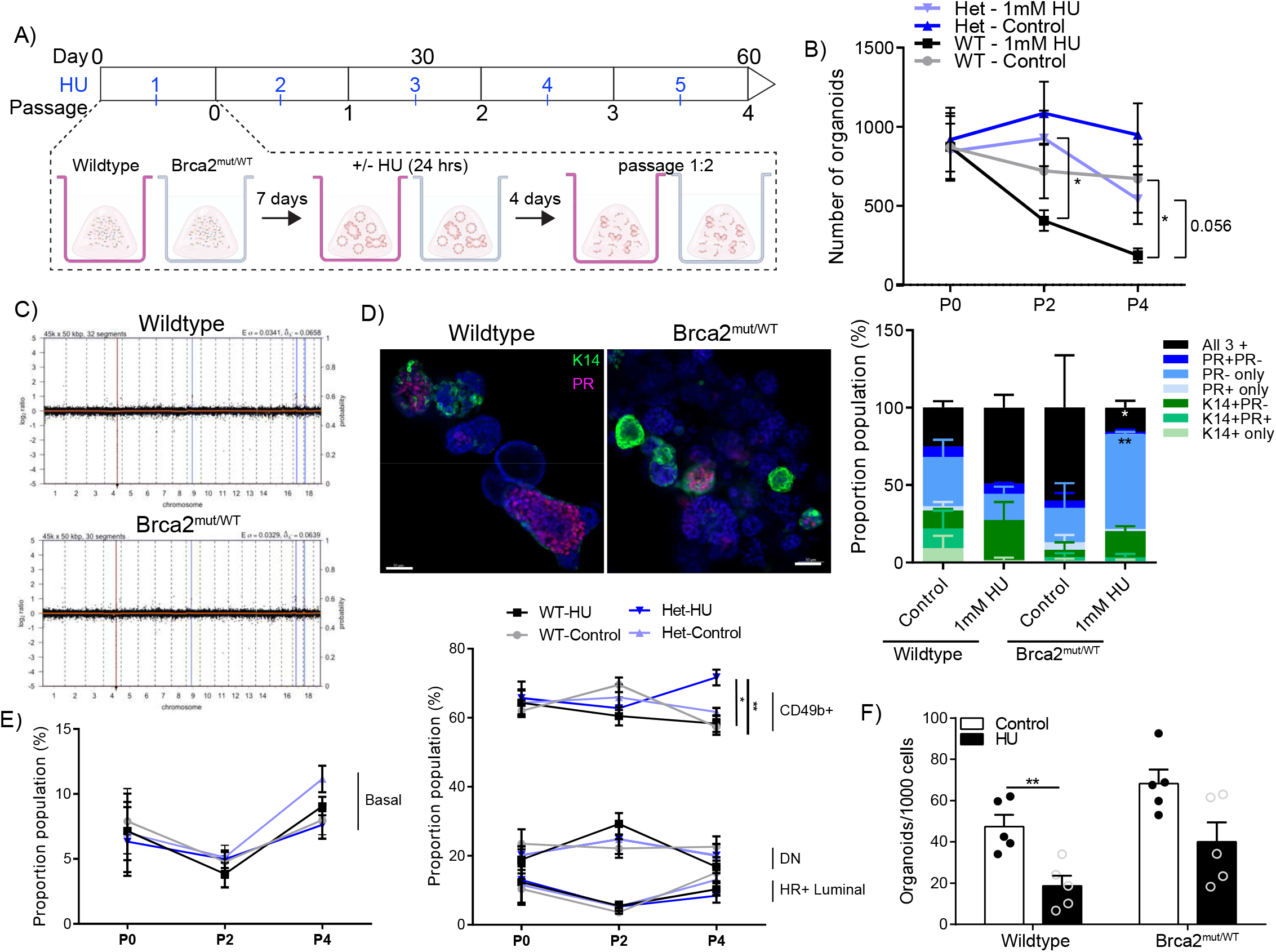
Intermittent genotoxic stress induces expansion of the HR- luminal population in Brca2^mut/WT^ organoids. A) Schematic illustrating the experimental pipeline for a longitudinal intermittent HU treatment of wildtype or Brca2^mut/WT^ mammary organoids. B) Quantification of the number of wildtype or Brca2^mut/WT^ organoids formed over multiple passages (P0, P2 and P4). Data presented as mean +/- SEM (n=5 per group). Mann-Whitney 2 tailed unpaired test was performed, *p=0.0278, n.s. p=0.0556. C) aCGH plots from wildtype (upper) or Brca2^mut/WT^ (lower) passage 4 HU treated organoids. Somatic CNAs were absent in HU treated organoids. Representative plots from 3 independent experiments are shown. D) Representative immunofluorescent images (left) of wildtype or Brca2^mut/WT^ organoids post HU treatment after passage 4. K14 (green), PR (magenta) and DAPI (blue). Scale bars, 50 µm. Quantification (right) of each organoid type at passage 4. Data presented as mean +/- SEM (n=3). An ANOVA followed by Tukey’s multiple comparison test was performed. Het HU all 3 + * p = 0.02. Het HU HR-only ** p = 0.006. E) Quantification of the percentage of basal (K14+, left) and luminal (K14-PR+ or K14-PR-right) cell types formed over multiple passages (P0, P2, P4) from wildtype or Brca2^mut/WT^ organoids treated with HU. Data presented as mean +/- SEM n=5-6. Mann-Whitney two-tailed unpaired t-test was used. * p = 0.013, ** p = 0.007. F) Quantification of the number of organoids formed from wildtype or Brca2^mut/WT^ single cells after 5 intermittent HU treatments. Data presented as mean +/- SEM (n=5). Student’s t-test was performed. ** p= 0.005.

Unchallenged wildtype and Brca2^mut/WT^ organoids retained their regenerative capacity and exhibited consistent organoid growth (Fig 3b). In contract, the number of wildtype HU-treated organoids diminished significantly (∼4-fold reduction, Fig 3b), indicative of reduced organoid regeneration. Unexpectedly, however, Brca2^mut/WT^ organoids exposed to repeated HU treatments retained regenerative capacity significantly better than HU-treated wildtype organoids. HU-exposed Brca2^mut/WT^ mammary organoids exhibited a statistically insignificant reduction in regenerative capacity compared with unchallenged Brca2^mut/WT^ controls by passage 4.Their regenerative capacity was >3-fold higher than that of similarly treated HU-exposed wildtype organoids (t-test, p 0.056) (Fig 3b). Thus, our results suggest that Brca2^mut/WT^ mammary organoids exhibit higher tolerance to replication stress than wildtype organoids.

Copy number alterations (CNAs) have recently been shown to accumulate in non-malignant cells carrying monoallelic *BRCA2* mutations^7^. To evaluate CNAs, we performed shallow whole genome sequencing (3x coverage) from unchallenged organoids at passage 0 and passage 4, and from HU-exposed organoids at passage 4, using 3 biological replicates. HU-treated Brca2^mut/WT^ organoids at passage 4 exhibited no increase in detectable CNAs, compared to wildtype organoids or untreated controls (Fig 3c, Supp Fig 3b-c). These findings suggest that HU-induced replication stress leads to alterations in the expansion of Brca2^mut/WT^ organoids without inducing chromosome instability marked by the accumulation of CNAs.

We next examined the cellular composition of HU-treated organoids. Wholemount staining for basal (K14) and HR+ luminal (Progesterone Receptor, PR) markers were carried out on 3 biological samples. Cells not expressing K14 or PR were categorised as HR- luminal. At passage 0 multilineage organoids (containing all 3 epithelial cell types) were present in 60%, 30%, 33% and 50% of wildtype, HU-treated wildtype (WT-HU), Brca2^mut/WT^, and HU-treated Brca2^mut/WT^ groups respectively (Supp Fig 3d). These trends continued in passage 2, however, HU-treated Brca2^mut/WT^ organoids showed a slight increase in HR- luminal cell proportion (Supp Fig 3e). Multilineage organoids were detected in 50%, 75% and 60% of wildtype, WT-HU and Brca2^mut/WT^ passage 4 organoids respectively (Fig 3d), demonstrating that HU treatments in wildtype mammary cells targeted all cell types equally. Notably, the HU-treated Brca2^mut/WT^ group significantly reduced the number of multilineage organoids to 30% with the remaining 65% containing only the HR- luminal cell type, compared with 10-40% in the other groups (Fig 3d). Our results indicate that prolonged HU treatments enabled the preferential survival and outgrowth of HR- luminal cells in Brca2^mut/WT^ organoids.

To examine the epithelial subtype proportions, we performed flow cytometry using our standard mammary antibody panels. In concordance with the wholemount findings, there were minimal changes detected in the proportions of basal and HR+ luminal cell types in any of the conditions tested (Fig 3e). In contrast, at passage 4 HU-treated Brca2^mut/WT^ organoids had a 20% increase in the proportion of HR- luminal cells compared to the other groups (Fig 3e). We next examined the organoid capacity from HU-treated cells. Passage 4 organoids were dissociated to single cells, reseeded into new organoid cultures and enumerated after 12 days. As expected, wildtype cell organoid formation capacity was severely reduced following prolonged HU treatments (Fig 3f). While Brca2^mut/WT^ HU treated cells displayed reduced organoid formation capacity, the proliferative ability was not as significantly impaired (Fig 3f). Taken together, we show that the Brca2^mut/WT^ HR- luminal population survives and expands following prolonged replication stress better than other luminal populations and better than wildtype cells.

### Single-cell profiling reveals HR- luminal cell expansion in Brca2^mut/WT^ mammary cells following replication stress exposure

We next sought to investigate the transcriptional changes associated with the enhanced Brca2^mut/WT^ organoid survival advantage following repeated replication stress. We performed scRNA-seq from early (passage 0) and late (passage 4) organoid culture time points. Across the four different conditions (wildtype or Brca2^mut/WT^ with or without HU treatments) and time points, a total of 34,200 cells were retained for analysis following quality controls. Unsupervised clustering across all high-quality cells and visualising the results using uniform manifold approximation and projections (UMAP) revealed 5 transcriptionally distinct subclusters (Fig 4a). Lineage specific markers were examined to identify the different clusters, and cell types were labelled using these marker genes. The dominant cluster was identified to be the HR- luminal population, which consisted of cells expressing known HR- luminal genes (*Lalba, Mfge8, Cd14, Elf5, cKit*), as well as a HR- luminal cycling population, which expressed HR- luminal genes, but also known cell cycle genes including *Birc5, Ccnb2, Cdc25c*, and *Cdkn2d*. A basal population was identified that expressed known basal markers (*Krt14, Id4, Pdpn, Tp63, Vim*), and a basal cycling population was also identified. The final cluster was identified to contain HR+ luminal cells expressing genes including *Areg, Prlr, Foxa1, Pgr*, and *Ar* (Fig 4b).

**Figure 4:**
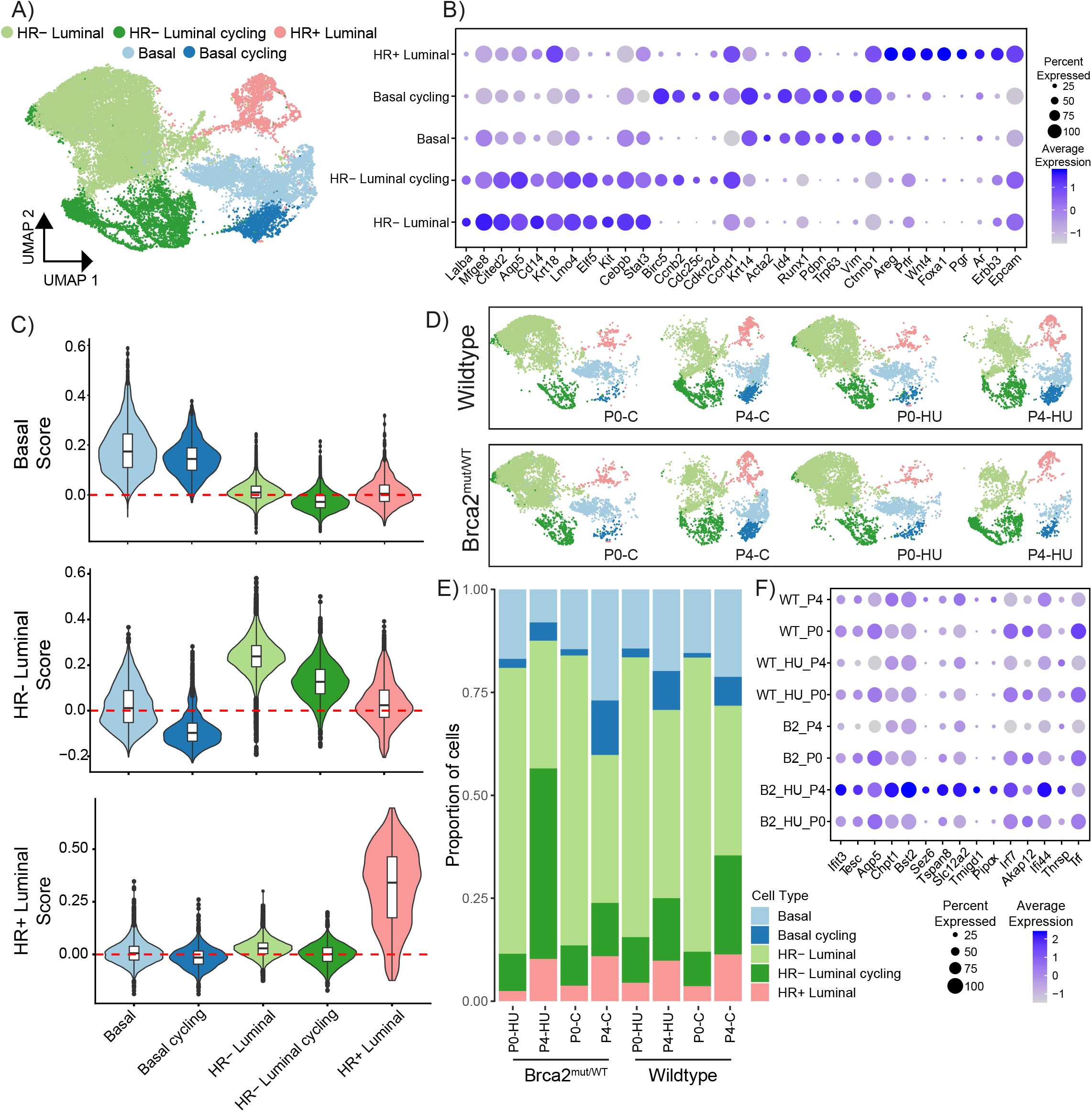
Single-cell transcriptomics reveal an expanded HR- luminal cycling population in Brca2^mut/WT^ organoids. A) Projection of dimensionality reduced (UMAP) scRNA-seq data (left, n = 34,200 cells in dataset from wildtype or Brca2^mut/WT^ organoids generated from 3 individual mice for each condition) coloured by cell type. B) Dot plot showing the expression of established mammary and proliferating cell markers associated with each cluster. Marker genes (x axis) for each major cell type cluster (y axis). Circle size indicates the percent of cells in the cluster expressing the gene. Filled colour represents the normalised and scaled mean expression level. C) Violin plots of Basal (upper), HR- Luminal (middle) and HR+ Luminal (bottom) cell scores across the mammary organoid cell clusters. UMAP analysis control (C) or HU treatment (HU) organoids at passage 0 (P0) or 4 (P4) wildtype or Brca2^mut/WT^ organoids; cell types are coloured. E) Frequency of cell types for each group, colour coded by cell type. F) Dot plot showing the expression of 15 of the top 30 differentially expressed marker genes in Brca2^mut/WT^ P4 HU treated organoids compared to the other groups. Circle size indicates the percentage of cells in which the gene expression was detected. Fill colour depicts the normalized and scaled mean expression level.

After annotating individual lineage-specific genes, we cross-examined mammary epithelial genes signatures derived from sorted HR-Luminal, HR+ Luminal and Basal cells^22^ to elucidate the extent to which organoids maintained lineage-specific programs. We scored each epithelial cluster from the organoids with respect to each lineage gene signature (Fig 4c). We found that mammary organoids faithfully recapitulated the gene expression signatures of primary mammary tissues. The Basal gene score was highest in the organoid basal and basal cycling clusters, the HR-Luminal score was highest in the organoid HR- luminal and HR- luminal cycling clusters, and the HR+ Luminal score was highest in the organoid HR+ luminal cluster (Fig 4c). UMAPs of wildtype and Brca2^mut/WT^ organoids resulted in all epithelial clusters being maintained (Fig 4d). Reassuringly, both wildtype and Brca2^mut/WT^ unchallenged and HU treated organoids at passage 0 and 4 showed the same gene signature correlations for the Basal score (Supp Fig 4a), HR-Luminal score (Supp Fig 4b), and the HR+ Luminal score (Supp Fig 4c). These findings demonstrate that mammary organoid cultures maintain the different lineages throughout the culture period irrespective of genotype or replication stress. Consistent with wholemount immunofluorescent analysis (Fig 3d), HU-treated Brca2^mut/WT^ organoids at passage 4 had the largest proportion of cells within the HR- luminal clusters, with the dominant population being the HR- luminal cycling cluster compared to Brca2^mut/WT^ unchallenged, or wildtype organoids (Fig 4e).

We then examined differences in gene transcription in Brca2^mut/WT^ HU treated cells compared to Brca2^mut/WT^ unchallenged and wildtype cells (+/-HU treatments) at passage 4. Among the most differentially expressed genes, interferon-related genes (*Ifit3, Irf7, Ifit44*) and luminal alveolar lineage-specific genes (*Tspan8, Thrsp, Aqp5*)^17,18,23^ were elevated in the Brca2^mut/WT^ HU treated group (Fig 4f). These genes were highly expressed in the Brca2^mut/WT^ HU treated passage 4 samples, and their expression increased over time (Fig 4f). Taken together, we showed that repeated stress events affect Brca2^mut/WT^ mammary epithelial cells differently. Basal cells reduced in number while HR- luminal cells significantly expanded. This expanded HR- luminal population also showed upregulation of transcripts.

### HR- luminal cells activate an interferon response after replication stress

To examine the impact of replication stress on the luminal compartment, we performed unsupervised clustering across the HR- luminal clusters, resulting in 9 transcriptionally distinct subclusters (Fig 5a-b). At passage 4, the HU-treated Brca2^mut/WT^ organoids had the highest proportion of cells in cluster 4 compared with Brca2^mut/WT^ unchallenged, or wildtype organoids (Supp Fig 5a). As cluster 4 was part of the HR- luminal cycling cluster, we next investigated the biological terms associated with this cluster. The GO biological terms for cluster 4 genes expressed in Brca2^mut/WT^ HU treated organoids were significantly attributed to Type 1 interferon responses (Fig 5c). Similarly, clusters 1, 2 and 3 in the Brca2^mut/WT^ HU treated organoids also showed enrichment in Type 1 interferon and neutrophil related response genes (Supp Fig 5b-d). Genes associated with these terms such as *Ifit3, Ir7, Ifit1, Blc2 and Stat1*, showed increased expression in the Brca2^mut/WT^ HU treated luminal cells compared with the other conditions, especially after prolonged exposures to replication stress (Fig 5d). Induction of immune-associated responses, including the genes we identified, have been reported in biallelic BRCA2 mutant cancer cells^24^. However, these interferon responses have not previously been demonstrated in mammary epithelial cells bearing monoallelic BRCA2 mutations. Our findings suggest that the expression of interferon response genes in Brca2^mut/WT^ HR- luminal cells is induced by persistent replication stress.

**Figure 5:**
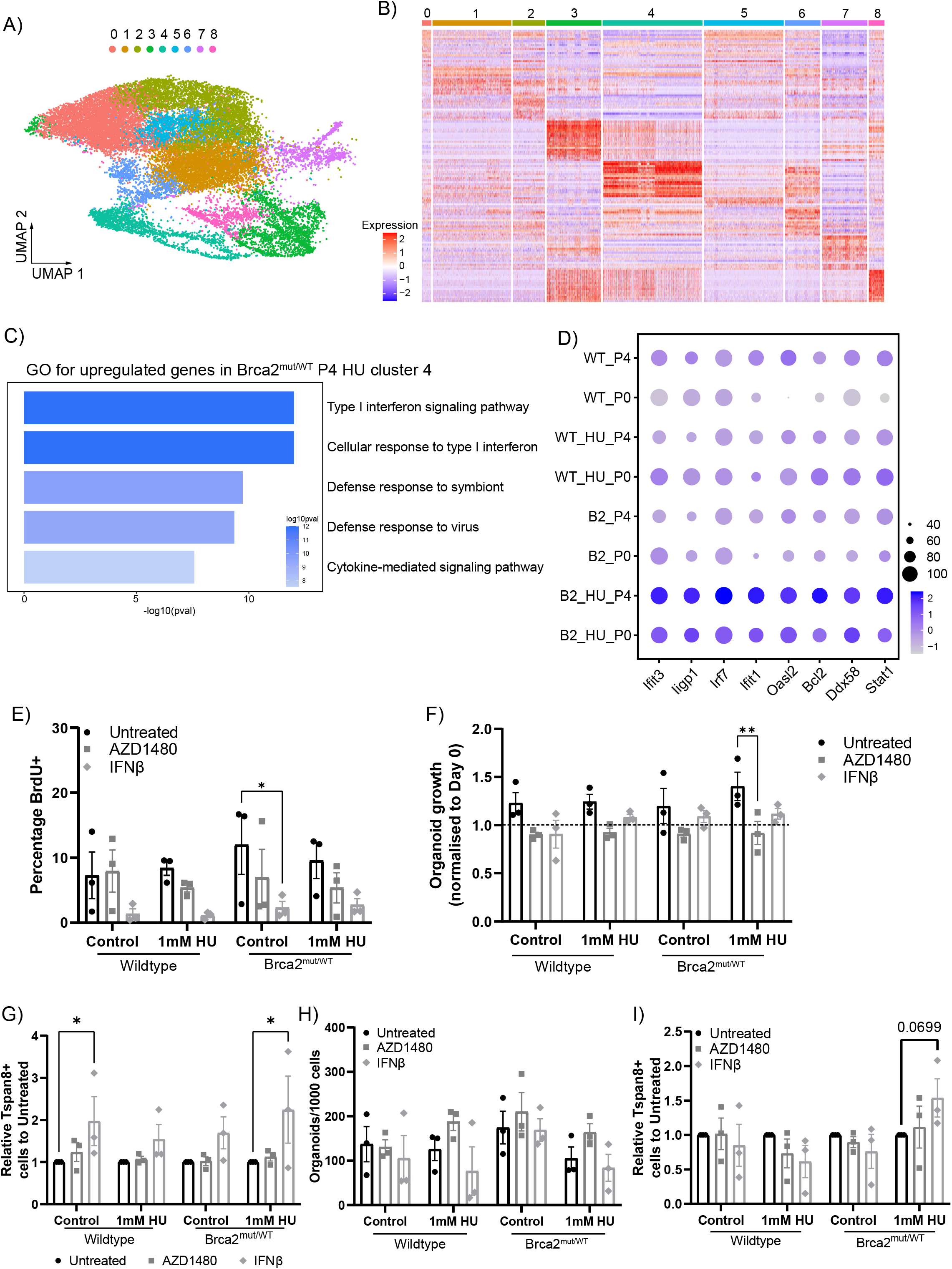
Investigating the HR- luminal cycle population shows increased Type I interferon response. Distribution of 9 HR- luminal epithelial cell subtypes on the UMAP. B) Heatmap of the normalized top 25 DEGs of HR- luminal clusters from all organoid groups. C) The top 5 GO Biological terms associated with the upregulated genes in Brca2^mut/WT^ HU passage 4 cluster 4 cells. D) Dot plot showing the expression of the genes identified in C) in Brca2^mut/WT^ P4 HU treated organoids compared to the other groups. Circle size indicates the percentage of cells in which the gene expression was detected. Fill colour depicts the normalized and scaled mean expression level. E) Quantification of the percentage of BrdU+ cells in the HR- luminal subpopulations from wildtype or Brca2^mut/WT^ passage 4 organoids after 4 days of IFNβ or AZD1480 treatment. Data presented as mean +/- SEM (n=3). An ANOVA followed by a Fishers LSD test was performed, *p=0.0141. F) Quantification of average organoid size after 4 days of INFβ or AZD1480 treatment normalised to Day 0 treatment organoid size. Dashed line indicates Day 0 area. Data presented as mean +/- SEM (n=3). An ANOVA followed by a Dunnett’s multiple comparison test, **p=0.0041. G) Quantification of the percentage of Tspan8+ cells in the HR- luminal population after 4 days of IFNβ or AZD1480 treatment on passage 4 wildtype or Brca2^mut/WT^ organoids. Data presented as the mean +/- SEM (n=3). An ANOVA followed by a Fishers LSD test *p=0.017. H) Quantification of the number of organoids formed from wildtype or Brca2^mut/WT^ passage 4 single cells after IFNβ or AZD1480 treatments. Data presented as mean +/- SEM (n=3). An ANOVA followed by a Fishers LSD test showed no statistical differences. I) Quantification of the percentage of Tspan8+ cells in the HR- luminal populations of wildtype or Brca2^mut/WT^ organoids from H). Data presented as the mean +/- SEM (n=3). An ANOVA followed by a Fishers LSD test showed no statistical differences.

To test the role of these interferon-related genes in organoid survival following replicative stress, we pre-treated passage 4 organoids for 3 days with recombinant Interferon β (IFNβ; inducing a type 1 interferon response) or a Janus Kinase inhibitor (Jaki; blocking Stat1 activation) before subjecting the organoids to HU treatments. While IFNβ treatment suppressed proliferation in all conditions (Fig 5e), it did not affect organoid size (Fig 5f), suggesting IFN signalling may provide an anti-apoptotic or other non-proliferative survival advantage. Conversely, Jaki did decrease organoid size specifically in HU-treated Brca2^mut/WT^ organoids (Fig 5f). These results suggest that the interferon response contributes to the survival advantage of Brca2^mut/WT^ organoids following replicative stress independently of proliferation.

The dominant cluster 4 population in the Brca2^mut/WT^ group is annotated as HR- luminal cycling population and is defined by high expression of mammary genes such as Tspan8 and Thrsp (Fig 4f). We reasoned Tspan8 could be utilised as a surrogate marker to identify this cluster. We observed an increase in Tspan8-positive luminal cells in all IFNβ treated organoids (Fig 5g). This increase was significantly higher in the Brca2^mut/WT^ HU treated and wildtype unchallenged luminal cells (Fig 5g). As expected IFNβ and Jaki treatments reduced proliferation and organoid growth following HU treatments, we next assessed whether HU treated cells were able to recover from these treatments and proliferate. To examine the proliferative potential after treatments, we reseeded single cells into new organoid cultures. In both IFNβ and Jaki treated groups, the cells recovered from growth inhibition to form comparable number of organoids to controls (Fig 5h). Flow assessment of the endpoint organoids demonstrated that increased Tspan8 expression was a transient phenotype following IFNβ withdrawal in wildtype and Brca2^mut/WT^ organoids, however, HU-treated Brca2^mut/WT^ organoids retained higher proportions of Tspan8+ cells (Fig 5i). In summary, replication stress induced a type I interferon response in Brca2^mut/WT^ HR- luminal cells prompting a cell survival response via entering a non-proliferative state for DNA repair and/or escape from cell toxicity. This response follows activation of the Jak pathway and increased Tspan8 expression allows a luminal cycling cell cluster to exist, enabling the HR- luminal cells to continue proliferation and growth.

### Deletion of HR- luminal cycling cluster genes compromise survival after replication stress

Our findings indicate that prolonged HU exposure caused an expansion of a HR- luminal cycling cluster. Overlay plots showed enrichment in the luminal cycling cluster 4 population, which contain high expression of *Tspan8* (Fig 6a-b). To validate whether Tspan8 positivity increases in Brca2^mut/WT^ HU treated cells, fresh mammary cells from 2 biological samples were seeded at passage 0, and the longitudinal assay was repeated. We observed Tspan8 expressing cells expanded in the HR- luminal cells of Brca2^mut/WT^ HU treated organoids, compared with unchallenged Brca2^mut/WT^ and wildtype (+/-HU treatment) organoids (Fig 6c). Next, we sought to identify whether Tspan8 was essential for Brca2^mut/WT^ organoids to sustain a growth advantage following replicative stresses. To genetically modify the organoids, we adopted the RNP CRISPR approach, where Cas9 protein and synthetic single stranded gRNA complexes were transiently introduced via nucleofection into the cells. This strategy has the advantage that the Cas9-gRNA complexes are rapidly degraded and minimises off-target effects compared to plasmid-based approaches^25^.

**Figure 6:**
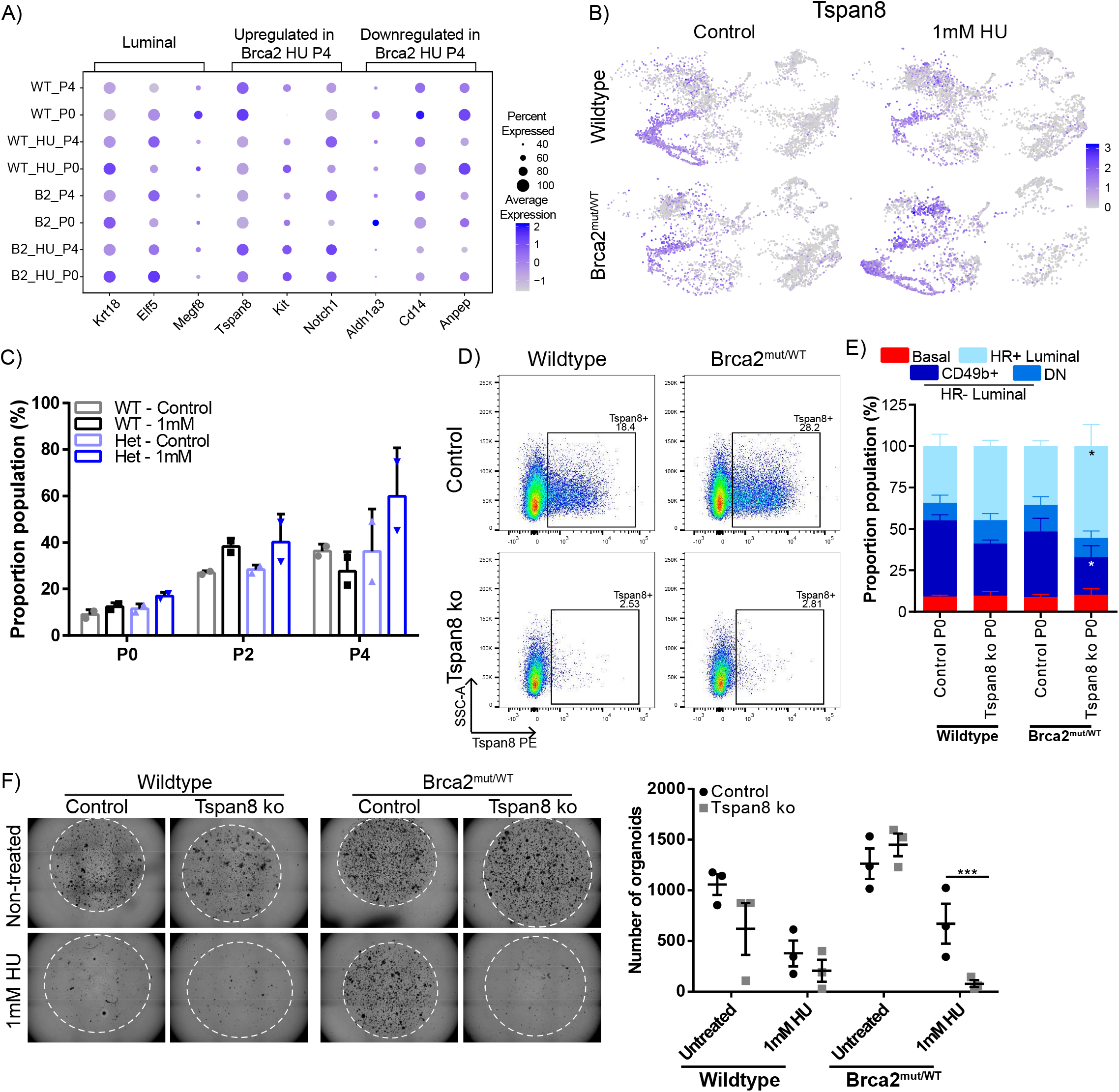
Deletion of Tspan8 expression in the HR- luminal cells render Brca2^mut/WT^ organoids incapable of recovery from DNA damage. A) Dot plot showing the expression selected HR- luminal expressed marker genes upregulated or downregulated in Brca2^mut/WT^ P4 HU treated organoids compared to the other groups. Circle size indicates the percentage of cells in which the gene expression was detected. Fill colour depicts the normalized and scaled mean expression level. B) UMAP with overlaid expression of Tspan8 gene from control or HU treated passage 4 wildtype or Brca2^mut/WT^ organoids. C) Quantification of the percentage of Tspan8+ cells in the luminal progenitor population over different passages (P0, P2 and P4) from wildtype or Brca2^mut/WT^ organoids treated with HU. Data presented as the mean +/- SD (n=2). D) Representative flow cytometry analysis of Tspan8 expression from control (upper) or Tspan8 knockout (ko, lower) wildtype and Brca2^mut/WT^ organoids. E) Quantification of the percentage of epithelial cell types from control or Tspan8 KO wildtype or Brca2^mut/WT^ organoids. Data presented mean +/- SD (n=3). An ANOVA followed by a Fishers LSD test, Het Tspan8 KO CD49b+ *p=0.0441, Het Tspan8 KO Sca1+ *p=0.0206. F) Representative brightfield images of Brca2^mut/WT^ organoids derived from control or Tspan8 KO cells at passage 3 after 4 intermittent HU treatments (left). Quantification (right) of the number of wildtype or Brca2^mut/WT^ organoids from control and Tspan8 KO post HU treatments at passage 3. Data presented mean +/- SEM (n=3). An ANOVA followed by a Tukey’s multiple comparison test, *** p<0.001.

Given that Tspan8 was highly expressed in cluster 4 (Fig 6b), we tested the role Tspan8 may play in HR- luminal survival and expansion under replication stress. We generated Tspan8 knockout (Tspan8 ko) mammary cells from primary 2D cultures and seeded cells into organoid cultures. Tspan8 ko editing efficiency was confirmed by DNA sequencing, and protein depletion was validated via flow cytometry analysis (Fig 6d). Tspan8 expression was detected in unedited wildtype and Brca2^mut/WT^ organoids at passage 3, as expected, while Tspan8 ko organoids maintained loss of Tspan8 expression at the beginning and after prolonged (1.5 months) organoid culture (Supp Fig 6a). Both wildtype and Brca2^mut/WT^ Tspan8 ko organoids exhibited a reduction in, but not complete absence of, the HR- luminal progenitor population (CD49b+ cells, Fig 6e). To determine whether Brca2^mut/WT^ organoids deficient of Tspan8 were able to survive replication stress, we repeated the longitudinal HU intermittent assay. The number of organoids formed during passage 3 from unchallenged Tspan8 ko wildtype or Brca2^mut/WT^ organoids were equivalent to non-edited (Fig 6f). As expected, prolonged replication stress reduced the number of wildtype organoids regardless of Tspan8 ko, indicating that wildtype organoids are less affected by Tspan8 loss (Fig 6f). Tspan8 ko almost completely eliminated the regenerative capacity of Brca2^mut/WT^ HU treated organoids (Fig 6f), reverting the replicative stress response akin to wildtype organoids.

To assess whether other genes expressed in cluster 4 contribute to this response we examined our scRNA-seq data and found that *Thrsp*, a gene expressed in luminal alveolar cells^17,18^, was also highly expressed in the HR- luminal cycling cluster 4 population (Supp Fig 6b). We generated Thrsp knockout (Thrsp ko) organoids following the RNP CRISPR pipeline and observed a consistent disruption of Thrsp via a 1 base pair out-of-frame insertion (Supp Fig 6c). Thrsp ko organoids phenocopied Tspan8 ko organoids in exhibiting a reduction in HR- luminal population in the Brca2^mut/WT^ organoids, with minimal impact on wildtype populations (Supp Fig 6d). The longitudinal HU intermittent exposure assay confirmed that Thrsp ko in wildtype organoids did not alter organoid growth dynamics, whereby similar number of organoids formed compared to non-edited organoids (Supp Fig 6e). Similar to the Tspan8 ko organoid response, Thrsp ko Brca2^mut/WT^ organoids were extremely sensitive to replication stress and significantly reduced organoid numbers (Supp Fig 6e). Taken collectively, our findings suggest that genetic deletion of Tspan8 or Thrsp genes, whose expressions are elevated in cluster 4, are required for maintenance of the HR- luminal cycling population, and that disruption of the HR- luminal cycling cluster 4 population significantly impairs the response to prolonged replication stress.

## Discussion

Cells heterozygous for pathogenic BRCA2 mutations do not apparently exhibit baseline phenotypes associated with loss of function in DNA repair. How heterozygous BRCA2 mutations affect epithelial cell lineages, which may predispose these cells towards the development of cancer, remains poorly understood. Here, we demonstrate that Brca2^mut/WT^ mammary organoids are not innately primed for aberrant behaviour, yet prolonged exposure to replication stress results in a preferential expansion of HR- luminal cells. This expanded population expressed genes characteristic of proliferation, type I interferon response and elevated mammary HR-alveolar genes (*Thrsp, Tspan8*) for sustained survival. Inactivation of these HR-alveolar genes reversed Brca2^mut/WT^ organoid survival response to replication stress.

Our data showing Brca2^mut/WT^ HR- luminal cell expansion raise important questions about the aetiology of BRCA2-associated breast cancers. While BRCA1 mutation carriers have an elevated risk for triple-negative breast cancers (TNBCs), BRCA2 mutation carriers develop breast cancers of different subtypes at a similar frequency to the general population and largely develop HR+ tumours^2,26^. Luminal HR+ tumours more closely resemble HR+ mammary epithelial cells^3,17,27^.

However, TNBCs are thought to arise from HR- luminal cells^4,28–30^, and premalignant changes identified in the breasts of BRCA1 carriers have mostly been in this cell type^3,31,32^. That we observe an expansion of HR- luminal cells after HU treatment in our organoids suggests our models may favour cellular advantage towards a HR-phenotype. Thus, the mechanisms of transformation giving rise to HR- and HR+ tumours in BRCA2 carriers may be distinct.

These distinct pathways preceding cellular transformation within the Brca2^mut/WT^ HR- luminal cells may be activated by replication-associated DNA damage responses. Activation of an intrinsic interferon response was observed in BRCA2-depleted breast cancer cells and has been attributed to endogenously arising DNA damage because these cells are defective in DNA repair^24,33^. However, our data indicates that the interferon response to replication stress may be a cell intrinsic phenotype in HR- luminal cells, which prompts them to enter a non-proliferative state that may promote DNA repair and/or escape from cell toxicity. This intrinsic interferon response includes a set of genes within the IFN-related DNA damage resistance signature (IRDS) (including **STAT1, IRF7, IFIT1/3, IFITM1**, ISG15, and **OAS** family, and **BST2** – those in bold are found in this study)^34^. Breast cancer cells with constitutive IRDS fail to transmit cytotoxic signals, resulting in pro-survival signals^34^, correlating with resistance to radiotherapy and chemotherapy in breast cancers^35^, but relatively little is known about how this pathway functions to facilitate the progression to tumorigenesis. Our data show increased expression of IRDS at the passage 4 time point, implicating that activated IRDS may drive a pro-survival signal in Brca2^mut/WT^ HR- luminal cells. Future studies may seek to elucidate the role of cell-intrinsic interferon signalling in aberrant cell formation and also determine the precise damage signal which activates this response.

In addition to the interferon response, we show that Brca2^mut/WT^ HR- luminal cells upregulate various of their lineage marker genes after repeated HU exposure and that *Tspan8* and *Thrsp* functionally enable a growth advantage in Brca2^mut/WT^ cells. In the mammary gland, high Tspan8 expression in myoepithelial cells identified a quiescent mammary stem cell phenotype, which became activated by ovarian hormones^36^. In this study, we observe increased Tspan8 positive cells, and upon depletion of Tspan8, HU-treated Brca2^mut/WT^ cells succumbed to replication stress, demonstrating both a dependence on *Tspan8* to stimulate the proliferative reservoirs necessary for cell survival and that *Tspan8* has a bona fide functional role in HR- luminal cells. Increased cell survival was also dependent on Thrsp expression. Thrsp stimulates fatty acid synthase (FASN)^37^, and increased levels of FASN have been shown to increase resistance to replicative drugs in vitro^38^. In our study, we also identified a novel interaction between interferon responses and the mammary HR- luminal cells lineage program (including Tspan8 and Thrsp). Future work to interrogate the expansion of HR- luminal cells in a prolonged replicative stress environment will determine the combined involvement of an interferon response and amplified lineage program in the emergence of transformative phenotypes.

Collectively, we establish a model for elucidating germline monoallelic Brca2 mutation cellular behaviour in different environmental conditions. DNA replication stress may arise from repeated cycles of hormone-driven expansion and remodelling during the mammary gland’s life cycle. In our model, unchallenged monoallelic Brca2 mutant mammary cells are not innately primed for survival behaviour. Only following multiple challenges do the HR- luminal Brca2^mut/WT^ cells exhibit a survival advantage by the combined and likely stepwise effects of IFN responses and *Tspan8*/*Thrsp* expression activating a luminal cycling population. Our data suggests that an expanded luminal population may identify Brca2 mutant cells that have activated a survival program and are adapted to endure stressful conditions.

## Methods

### Mice

Mammary glands from Brca2 germline and wildtype mouse strains were provided by the Venkitaraman group. Tissues collected from virgin adult young (3-month-old) and aged (7–10-month-old) female Brca2 Tr/WT^19^ and littermate Brca2 WT/WT mice. All mice were treated in strict accordance with the local ethical committee (University of Cambridge Licence Review Committee) and the UK Home Office guidelines. Animals were housed in a pathogen-free environment with ab libitum to diet and sanitised water. Mice were housed in a standard facility and maintained under standard conditions: 20-24°C, 40-70% humidity and a 12h light/dark cycle.

### Organoids cultures

#### Mammary organoids

Mammary epithelial cells were collected from third and fourth mammary glands of virgin mice. Briefly, lymph nodes were removed, and glands minced with surgical scissors before enzymatical dissociation for 1.5h in DMEM/F12 (1:1) supplemented with 2 mg mL^−1^ collagenase (Roche) + Gentamicin (Gibco). Samples were briefly vortexed every 30min. Mammary gland fragments were treated with NH_4_CL to lyse red blood cells, then dissected to single cells with 0.05% Trypsin-EDTA (STEMCELL Technologies) and 5 mg ml^-1^ dispase (STEMCELL Technologies) and 1 mg ml^-1^ DNAse (Sigma) and filtered through a 40 µm cell strainer (Falcon).

Single cells were mixed with Matrigel (Corning) and seeded (5500 cells per 35 µl Matrigel) in 24-well plates. In all, 500 µl of MammoCult™ Organoid Growth Medium (Mouse, STEMCELL Technologies) + Gentamicin (Gibco) was added per well and medium was replenished every other day. Cultures were maintained in a 37°C humidified atmosphere under 5% CO_2_ for 10-12 days. For replicative stress experiments: Hydroxyurea (1 mM, unless otherwise indicated, Sigma) was added to organoid cultures on day 5/6 for 24h. Cultures washed with PBS and replenished with fresh medium. For inhibitor experiments: Recombinant Interferon β (250U/ml, Peprotech) or Jak inhibitor (AZD1480 100nM; Calcembio?0) was added to passage 4 organoids on day 5 for a total of 4 days. Replicative stress was performed on day 7 by administering 1mM HU to the organoids for 24 h.

#### Organoid passaging

Mammary organoids were maintained in culture by passaging them every 10-12 days at a 1:2 ratio split. Briefly, mammary organoids were washed by PBS and mechanically released from the Matrigel by breaking the matrix with a P1000 pipette and treated with TrypLE (Invitrogen) for 2-3 min at 37 °C to generate fragments. Following washes with trypsin inhibitor, mammary fragments were centrifugated at 100g for 5min at 4°C. Supernatant containing single cells was removed and the fragments were resuspended in Matrigel, seeded in 24-well plates, and exposed to previously described culture conditions. For quantitative organoid formation analysis: organoids were mechanical dissociated in TrypLE for 10-15 minutes at 37 °C to generate a single cell suspension.

Single cells were resuspended in Matrigel and cultured. Organoids were imaged on a SX5 Incucyte® (Sartorius). Organoids were imaged at the start and end of every passage. For inhibitor experiments: organoids were imaged at the start and end of treatments. Organoid numbers were calculated using the Incucyte® organoid analysis software module (Sartorius) CRISPR edited cells: 4000 single cells were mixed with 35 µl Matrigel in 24-well plates and supplemented with 500 µl of MammoCult™ Organoid Growth Medium (Mouse) and cultured as previously described.

#### Organoid cryopreservation

At times, organoids were grown for the complete 10-12 days, before removal from Matrigel by breaking the matrix with ice-cold DMEM followed by a PBS wash. Organoids were resuspended in freezing media (Cryostor 10, Sigma) at a density equivalent to 2 wells per ml freezing solution, and aliquoted into cryovials. Cryovials were stored overnight at -80°C before long term storage in liquid nitrogen. For thawing, vials were placed in placed in a 37°C water bath and the contents washed twice in DMEM + 2% FBS (Invitrogen), before reseeding in Matrigel at required density.

### CRISPR

#### Cell preparation

Mammary glands from 8–10-month-old virgin Brca2 Tr/WT and Brca2 WT/WT were processed to single cells as described above and seeded into 6-well plates at a density of 1-2 x10^5^ cells/well. EpiCult™ Plus Medium (STEMCELL tech) + 50 mg ml^-1^ Hydrocortisone (Sigma) + Gentamicin (Gibco) was added to the wells. Cultures were maintained in a 37°C humidified atmosphere under 5% CO_2_ for 8-10 days. Medium change every 3 days. Cells were trypsinsed at 70-80% confluency, using pre-warmed 0.05% Trypsin at 37°C for 5 minutes. The reaction was terminated by adding DMEM + 2% FBS. Cells were passed through a 40 µm cell strainer.

#### Nucleofection for gene targeting

1 µl Cas9 protein (5 µg/µl, Thermo Fisher) and 100 pM of synthetic guide RNA (Tspan8 guide sequence 5’ – 3’: GGGGAGTTCCGTTTACCCAA; Thrsp guide sequence 5’ – 3’: AGTCATGGATCGGTACTCCG; Merck) were mixed and incubated at RT for a minimum of 10min to assemble the ribonucleoprotein (RNP) complex. 2 × 10^5^ single cells were resuspended with Lonza P3 Primary Cell Nucleofector® Solution and mixed with the pre-formed RNP complexes and transferred to a Nucleocuvette™ (Lonza). Nucleofection was performed using program EO-115 on the 4D Nucleofector™ X unit (Lonza). After nucleofection, the cells were immediately transferred back to warm EpiCult™ Plus complete medium to continue culture for a further 5-7 days.

#### Genotyping

Cells were harvested 5-7 days after nucleofection, and genomic DNA was extracted with the DNeasy Blood & Tissue Kit (Qiagen). Targeted regions were PCR amplified using Q5® High-Fidelity DNA Polymerase (NEB), with the corresponding primers listed. Tpan8 forward primer: AAGACACATCTCCGTAACGACA, reverse primer: AGCTCCCCTGGTGCTTACTG; Thrsp forward primer: CGGACTCTGAGGAAGGAAGC, reverse primer: GGTGGAACTGGGCTTCTAGG. The products were gel purified using QIAquick PCR Purification Kit (Qiagen) and sent for sanger sequencing.

### Immunofluorescent staining

Wholemounts: Organoids were washed with 1x PBS and mixed with 1 mL of Cell Recovery Solution. The mixture was incubated on ice for 40min, centrifuged at 300g, then washed twice with 1x PBS. Organoids were fixed with 4 % paraformaldehyde for 45min at RT, washed 3x with PBS, 0.1% BSA, 0.2% Triton X-100 and 0.1% Tween 20 (Immunofluorescence (IF) Buffer) and then heat-induced antigen retrieval in citrate buffer (pH 6) for 20min. Organoids were then permeabilised with 1% Triton X-100 in PBS for 1h and blocked with 5% goat serum in IF buffer for 1h. Organoids were incubated on a shaker for 24-48h at RT with primary antibodies in IF buffer (Table S1). After incubation with the primary antibodies, organoids were washed three times with IF buffer for 5min and incubated with the secondary antibodies diluted in IF buffer supplemented with 10% goat serum (Table S2) and incubated on a shaker for 24-48h. Nuclei were counterstained with DAPI. Organoids were cleared using fructose glycerol^39^. Organoids were imaged using the Zeiss 880 confocal microscope (Zeiss).

Tissue sections: Intact mammary glands were freshly isolated and fixed in 10% neutral buffered formalin overnight before processing the tissue into paraffin. Tissue blocks were sectioned at 4 μm, deparaffinized and before performing heat-induced antigen retrieval in citrate buffer (pH 6). The samples were preblocked in PBS with 1% BSA and 0.1% Tween 20, then incubated with primary antibodies overnight at 4 °C (Table S1). The secondary antibodies were goat anti-AF488, goat anti-Rabbit Cy3, and goat anti-AF647 (Table S2; Jackson ImmunoResearch) and were all used at 2 μg ml^−1^. A no primary antibody was used as a control. Slides were stained with DAPI to visualize the nuclei and sections mounted with ProLong Gold antifade (Invitrogen). Tissue sections were imaged using the Zeiss 880 confocal microscope (Zeiss).

### Mass Cytometry sample preparation and analysis

Mass cytometry and subsequent data analyses were performed as previously reported^17,40^. In brief, samples were barcoded using the Cell-ID 20-Plex Pd Barcoding Kit (Fluidigm) following manufacturer’s instructions. Samples were then pooled, resuspended in HBSS (Gibco) with 100 µg/mL DNAse I (STEMCELL Technologies) for 15 min in a 37°C water bath, and washed once with Cell-Staining Medium (CSM; Fluidigm). Next, the pooled sample was stained at room temperature for 30 minutes with an antibody cocktail containing the appropriate volume of each antibody with CSM. The sample was washed two times in CSM, fixed for 30 min at room temperature in 4% PFA (EMS) in PBS, and washed once with CSM. The sample was then suspended in 0.5 mL Intercalator Solution (300 μL CSM with 50 μL 16% PFA, 50 μL Fix-Perm Buffer (Fluidigm), and 0.67 μL Cell-ID Intercalator-Ir (Fluidigm)) per 1 million cells overnight at 4°C. The sample was washed once the following day with CSM and twice with Milli-Q water. The sample was then resuspended in bead water (1:10 4-Element EQ Beads (Fluidigm) in Milli-Q water) to a concentration of ∼750,000 cells/mL solution. Samples were run using the Helios Mass Cytometer.

CyTOF data were normalized to the bead signal, converted to an FCS format, and debarcoded using the CyTOF Software v7 from Fluidigm. Samples were gated in FlowJo for quality control parameters to yield live, single cells: Event Length, Gaussian Parameters, 140Ce-Beads, DNA content 191Ir, DNA content 193Ir, and Viability 195Pt. All single cells were then imported into the toolkit Scanpy^41^, embedded as a UMAP, and clustered using the Leiden algorithm^42^ to perform lineage gating. Heatmaps of the CyTOF data were made in R using the heatmap.plus, RColorBrewer, and gplots packages.

### Flow cytometry

Mammary glands: Dissociated to single cells and cells were then incubated with the following primary antibodies (Table S3): CD31, CD45, Ter119, EpCAM, CD49f,CD49b, and Sca1. Biotin conjugated antibodies were detected with Streptavidin-eFluor450 (eBioscience). Cells were then filtered through a 30-μm cell strainer (Partec) and incubated with DAPI and were sorted on a FACSAria II (Becton Dickinson). Organoids: Dissociated to single cells using TrypLE at 30°C for 10-15mins. Single mammary cells were then incubated with EpCAM, CD49f, CD49b, Sca1, and where required Tspan8. Cells were then filtered through a 30-μm cell strainer and incubated with DAPI and were analysed by using an LSRII (Becton Dickinson). Flow cytometry data were analysed using FlowJo (version 10. Tree StarInc. For BrdU-staining: BrdU (100uM) was administered to organoids for 12h then digested to single cells as described above. For intracellular staining, cells were first stained with the indicated surface markers and then fixed with BD Cytofix/Cytoperm Buffer (BD Bioscience) for 20 min at 4 °C, followed by incubation with BD Cytoperm Plus Buffer for 10 min at 4 °C and re-fixed for 5 min at 4 °C. Cells were then treated with DNase I (1 mg/ml, Sigma) in PBS and then immunostained with BrdU-FITC. For γ-H2AX staining: cells were stained with Zombie UV (BioLegend) instead of DAPI and then fixed with BD Cytofix/Cytoperm Buffer (BD Bioscience) for 20 min at 4 °C, followed by incubation with BD Cytoperm Plus Buffer for 10 min at 4 °C and re-fixed for 5 min at 4 °C. Cells were immunostained with Phospho-Histone H2A.X (Ser139)-FITC on ice for 30 min.

### Single cell RNA sequencing

#### Cell preparation

For single cell RNAseq: cryopreserved organoids from P0, P2 and P4 were thawed, reseeded into Matrigel, and cultured for 3 days. Organoids were dissociated into single cell suspension as described above. Viability of <90% was confirmed on all samples and cells were immediately submitted to the CRUK CI genomics core facility for library preparation.

#### Library preparation and sequencing

Library preparation was performed according to instruction in the 10× Chromium single-cell kit version 3. The libraries were then pooled and sequenced across eight lanes on a NovaSeq6000 S2.

#### Bioinformatics analysis of scRNA-seq data

Cell Ranger 6.0.2 (http://10xgenomics.com) was used to process Chromium scRNA-seq output and generate the count table. Samples were demultiplexed using barcode assignment and unique molecular identifier (UMI) quantification. FASTQ reads were aligned to the mouse reference genome refdata-cell ranger-mm10-3.0.0. Seurat (V4.0) in R (V4.1.3) was used to carry out all analyses. Cells that met quality control conditions (unique number of genes between 1000 and 6000 and <5% of genes are mitochondrial genes per cell) were included for downstream analysis. Thirty-four thousand two hundred cells with expression levels for 3000 genes passed quality control. Individual samples were integrated, data normalized and scaled with the Sctransform function. The dimensional reduction was performed by PCA. The FindNeighbors function of Seurat was used to construct the Shared Nearest Neighbor (SNN) Graph, based on unsupervised clustering performed with Seurat function FindClusters. For visualisation, the dimensionality was further reduced using Uniform Manifold Approximation and Projection (UMAP) method with Seurat function RunUMAP. Differentially expressed genes were determined using Wilcoxon rank sum test with P-value < 0.05.

We determined the major epithelial cell subsets whereby differentially expressed genes (DEGs) of each cluster were identified using the FindMarkers function in Seurat, which returns the gene names, average log fold-change, and adjusted p-value for genes enriched in each cluster. We carefully reviewed top 50 DEGs for each cluster with special focus on well-studied mouse mammary epithelial markers. These were integrated to define cell types and cell transcriptomic states.

Mammary primary epithelial cell signatures were curated from several previously published mammary scRNA-seq datasets^22^. The gene signatures of basal, HR- Luminal and HR+ Luminal mammary primary cells was then calculated between the clusters in our data with the AddModuleScore function in Seurat. Heatmaps of the HR- luminal cell clusters were plotted using the doHeatmap Seurat function using the top 25 genes from each luminal cluster. Gene Ontology enrichment analysis of DEGs across the clusters and conditions was performed using the DEenrichRPlot function in Seurat. The EnrichR database used was the GO_Biological_Process_2021.

### Whole genome sequencing

#### Low pass whole genome sequencing

Shallow whole genome sequencing was performed at Novogene’s Cambridge Sequencing Centre. Passage 0 and 4 organoids were dissociated to single cell suspensions and DNA extracted using the QIAamp DNA Micro kit (Qiagen). The extracted genomic DNA was sheared and the fragments were end repaired, A-tailed and further ligated with Illumina adapters. The fragments with adapters were PCR amplified, size selected, and purified. The library was checked with Qubit and real-time PCR for quantification and bioanalyzer for size distribution detection. The 150 bp paired-end sequencing reaction was performed, resulting in an average genome coverage of 3× per sample.

#### CNA analysis

Raw reads were trimmed for sequencing adaptors with fastp using default parameters. Trimmed reads were then aligned to the reference genome using Burrows-Wheeler Aligner (BWA). Subsequent processing, including duplicate removal was performed using samtools and Picard. Alignment statistics and genome coverage metrics were extracted using GATK and Picard.

CNA calling and analysis was performed using QDNAseq package in R. Bin annotations for mouse genome build were obtained from QDNAseq.mm10 package. Segmentation analysis was performed using default parameters and the resultant output files were summarised using R.

## Statistics

Data are presented as mean ± SD or mean± SEM, as indicated in the figure legends. All data was obtained from at least two independent biological replicates and n represents the number of independent mice or samples used for the analysis. N values are indicated in the figure legends. Statistical significance was determined with two-sided Mann–Whitney test or t-test with Welch correction and reported from GraphPad Prism v9. Differences were significant when P < 0.05. Differences around P<0.05 were listed in the figure legends.

## Acknowledgements

We thank Dr Ben Hall (UCL) and Dr Harveer Dev (Department of Oncology, Cambridge) for critical reading of our manuscript. We thank the CIMR flow cytometry facility for assistance with cell sorting and the CRUK Cambridge Institute genomics core facility, in particular Katarzyna Kania with preparing the samples for single cell RNA-sequencing, and the DFCI Mass Cytometry Core, led by Nicole Paul and Eric Haas. This work was funded by Gray Foundation Team Science Award to JB and ARV, Medical Research Council (MRC) Programme grants MC_UU_12022/1 and MC_UU_12022/8 to A.R.V., and by the Krishnan-Ang Fellowship to M.S.

## Author information

### Contributions

MS and ARV conceived this study. MGN, GKG, LRK, KG, DP and MS performed experiments and analysed experimental data. MS, JB and ARV interpreted the results. MS and ARV wrote the paper, which was edited by all of the authors.

## Ethics declarations

All authors declare no competing interests

## Supplemental figures

Supplemental Figure 1. Signal intensities of A) PR, B) CD36 and CD14, and C) E-Cadherin. All epithelial ducts (60-210) from a whole mammary gland were imaged for each antibody and mean fluorescent intensities values reported. Data presented mean +/-SD (n=3). Kruskal-Wallis followed by Dunn’s multiple comparison test was performed. A) **p=0.004, B) CD36 *p= 0.023, CD14 *p=0.0109, C) E-Cadherin ***p=0.0005, **p=0.005. D) Schematic illustrating the experimental pipeline of culturing mammary epithelial organoids. E) Brightfield timeline images of mammary cells detailing propagation of organoid development after seeding as non-sorted single cells. Representative images from three independent experiments. F) Representative immunofluorescent images of mammary gland organoids after 12 days culturing. K14 (green), PR (magenta). Representative images on the left and E-Cadherin (green) and Vimentin (magenta) representative images on the right. Nuclear staining with DAPI (blue). Representative images from three independent experiments. Scale bars, 50 µm. G) Brightfield images of Basal, HR- luminal and HR+ luminal flow sorted cells after seeding (P0) and imaged daily. Representative images from three independent experiments. H) Representative immunofluorescent images of mammary gland organoids derived from Basal (left), HR- luminal (middle) or HR+ luminal cells after 12 days culturing. K14 (green), PR (magenta), nuclear staining with DAPI (blue). Representative images from three independent experiments. Scale bars, 50 µm.

Supplemental Figure 2. A) Quantification of the percentage of γH2AX+ cells from wildtype or Brca2^mut/WT^ organoids after 4mM HU treatment. Data depicts the different epithelial populations. Data presented as the mean +/- SEM (n=5-6). B) Quantification of the proportion of epithelial populations in the wildtype or Brca2^mut/WT^ organoids after 4mM HU treatment. Data presented as mean +/- SEM (n=5-6). C) Quantification of the percentage of BrdU+ cells in the different epithelial subpopulations from wildtype or Brca2^mut/WT^ organoids after 4mM HU treatment. Data presented as mean +/- SEM (n=5-6). An ANOVA followed by Fishers LSD showed no significant differences.

Supplemental Figure 3. A) Quantification of the percentage of γH2AX+ cells from wildtype or Brca2^mut/WT^ organoids after 1mM HU treatment over multiple passages (P0, P2 and P4). Data depicts the different epithelial populations. Data presented as the mean +/- SEM (n=3). B) Bar chart of the total CNVs detected of sWGS in wildtype or Brca2^mut/WT^ organoids from passage 0 and passage 4. Data presented as mean +/- SEM (n=3). C) aCGH plots from wildtype (upper) or Brca2^mut/WT^ (lower) organoids at passage 0 and passage 4. Somatic CNAs were absent in all organoids. Representative plots from 3 independent experiments are shown. (D-E) Quantification of immunofluorescent images of wildtype or Brca2^mut/WT^ organoids post HU treatment stained with K14 (green), PR (magenta) and DAPI (blue) at D) passage 0 or E) passage 2. Data presented as mean +/- SD (n=3). An ANOVA followed by Tukey’s multiple comparison test showed no statistical differences.

Supplemental Figure 4. Violin plots of A) Basal, B) HR- Luminal and C) HR+ Luminal cell scores across the mammary cell clusters of wildtype or Brca2^mut/WT^ unchallenged and HU treated organoids from passage 0 or 4.

Supplemental Figure 5. A) Frequency of HR- luminal cell clusters for each group, colour coded by cluster. The top 5 GO Biological terms associated with the upregulated genes in Brca2^mut/WT^ HU passage 4 of B) Cluster 1, C) Cluster 2 and D) Cluster 3 cells.

Supplemental Figure 6. A) Representative flow cytometry analysis of Tspan8 expression from control (upper) or Tspan8 knockout (ko, lower) wildtype and Brca2^mut/WT^ organoids at passage 0 and passage 3. B) UMAP with overlaid expression of *Thrsp* gene from control or HU treated passage 4 wildtype or Brca2^mut/WT^ organoids. C) Illustration of Sanger sequencing results for the *Thrsp* indel created by CRISPR/Cas9 in wildtype and Brca2^mut/WT^ mammary cells. D) Quantification of the percentage of epithelial cell types from control or Thrsp ko wildtype or Brca2^mut/WT^ organoids. Data presented mean +/- SD (n=2/3). An ANOVA followed by a Fishers LSD test, *p=0.0225, **p=0.0067. E) Quantification of the number of wildtype or Brca2^mut/WT^ organoids from control and Thrsp ko post HU treatments at passage 3. Data presented mean +/- SEM (n=3). An ANOVA followed by a Fishers LSD test, *p=0.0463.

## References

1. Roy, R., Chun, J. & Powell, S. N. BRCA1 and BRCA2: different roles in a common pathway of genome protection. Nat Rev Cancer 12, 68–78 (2011).

2. Larsen, M. J. et al.. Classifications within molecular subtypes enables identification of BRCA1/BRCA2 mutation carriers by RNA tumor profiling. PLoS One 8, (2013).

3. Lim, E. et al.. Aberrant luminal progenitors as the candidate target population for basal tumor development in BRCA1 mutation carriers. Nat Med 15, 907–913 (2009).

4. Molyneux, G. et al.. BRCA1 basal-like breast cancers originate from luminal epithelial progenitors and not from basal stem cells. Cell Stem Cell 7, 403–417 (2010).

5. Casey, A. E. et al.. Mammary molecular portraits reveal lineage-specific features and progenitor cell vulnerabilities. Journal of Cell Biology 217, 2951–2974 (2018).

6. Ding, L. et al.. Perturbed myoepithelial cell differentiation in BRCA mutation carriers and in ductal carcinoma in situ. Nat Commun 10, (2019).

7. Karaayvaz-Yildirim, M. et al.. Aneuploidy and a deregulated DNA damage response suggest haploinsufficiency in breast tissues of BRCA2 mutation carriers. Sci Adv 6, (2020).

8. Choudhury, S. et al.. Molecular profiling of human mammary gland links breast cancer risk to a p27+ cell population with progenitor characteristics. Cell Stem Cell 13, 117–130 (2013).

9. Cheung, A. M. Y. et al.. Brca2 deficiency does not impair mammary epithelium development but promotes mammary adenocarcinoma formation in p53(+/-) mutant mice. Cancer Res 64, 1959–1965 (2004).

10. Kass, E. M., Lim, P. X., Helgadottir, H. R., Moynahan, M. E. & Jasin, M. Robust homology-directed repair within mouse mammary tissue is not specifically affected by Brca2 mutation. Nat Commun 7, (2016).

11. Patel, K. J. et al.. Involvement of Brca2 in DNA repair. Mol Cell 1, 347–357 (1998).

12. Tutt, A. et al.. Absence of Brca2 causes genome instability by chromosome breakage and loss associated with centrosome amplification. Curr Biol 9, 1107–1110 (1999).

13. Moynahan, M. E., Pierce, A. J. & Jasin, M. BRCA2 is required for homology-directed repair of chromosomal breaks. Mol Cell 7, 263–272 (2001).

14. Joshi, P. A. et al.. Progesterone induces adult mammary stem cell expansion. Nature 465, 803–807 (2010).

15. Giraddi, R. R. et al.. Stem and progenitor cell division kinetics during postnatal mouse mammary gland development. Nat Commun 6, (2015).

16. Shehata, M. et al.. Proliferative heterogeneity of murine epithelial cells in the adult mammary gland. Commun Biol 1, (2018).

17. Gray, G. K. et al.. A human breast atlas integrating single-cell proteomics and transcriptomics. Dev Cell 57, 1400-1420.e7 (2022).

18. Bach, K. et al.. Differentiation dynamics of mammary epithelial cells revealed by single-cell RNA sequencing. Nat Commun 8, (2017).

19. Friedman, L. S. et al.. Thymic lymphomas in mice with a truncating mutation in Brca2. Cancer Res 58, 1338–1343 (1998).

20. Nguyen-Ngoc, K. V. et al.. 3D culture assays of murine mammary branching morphogenesis and epithelial invasion. Methods Mol Biol 1189, 135–162 (2015).

21. Flurkey, K., M. Currer, J. & Harrison, D. E. Chapter 20 - Mouse Models in Aging Research. in The Mouse in Biomedical Research (Second Edition) (eds. Fox, J. G.et al.) 637–672 (Academic Press, 2007).

22. Saeki, K. et al.. Mammary cell gene expression atlas links epithelial cell remodeling events to breast carcinogenesis. Commun Biol 4, (2021).

23. Li, C. M. C. et al.. Aging-Associated Alterations in Mammary Epithelia and Stroma Revealed by Single-Cell RNA Sequencing. Cell Rep 33, (2020).

24. Reisländer, T. et al.. BRCA2 abrogation triggers innate immune responses potentiated by treatment with PARP inhibitors. Nat Commun 10, (2019).

25. Sun, D. et al.. A functional genetic toolbox for human tissue-derived organoids. Elife 10, (2021).

26. Ha, S. M. et al.. Association of BRCA Mutation Types, Imaging Features, and Pathologic Findings in Patients With Breast Cancer With BRCA1 and BRCA2 Mutations. AJR Am J Roentgenol 209, 920–928 (2017).

27. Shehata, M. et al.. Phenotypic and functional characterisation of the luminal cell hierarchy of the mammary gland. Breast Cancer Res 14, (2012).

28. Gusterson, B. A., Ross, D. T., Heath, V. J. & Stein, T. Basal cytokeratins and their relationship to the cellular origin and functional classification of breast cancer. Breast Cancer Res 7, 143–148 (2005).

29. Smart, C. E. et al.. Analysis of Brca1-deficient mouse mammary glands reveals reciprocal regulation of Brca1 and c-kit. Oncogene 30, 1597–1607 (2011).

30. Bach, K. et al.. Time-resolved single-cell analysis of Brca1 associated mammary tumourigenesis reveals aberrant differentiation of luminal progenitors. Nat Commun 12, (2021).

31. Nolan, E. et al.. RANK ligand as a potential target for breast cancer prevention in BRCA1-mutation carriers. Nat Med 22, 933–939 (2016).

32. Sau, A. et al.. Persistent Activation of NF-κB in BRCA1-Deficient Mammary Progenitors Drives Aberrant Proliferation and Accumulation of DNA Damage. Cell Stem Cell 19, 52–65 (2016).

33. Xu, H. et al.. Up-regulation of the interferon-related genes in BRCA2 knockout epithelial cells. J Pathol 234, 386–397 (2014).

34. Weichselbaum, R. R. et al.. An interferon-related gene signature for DNA damage resistance is a predictive marker for chemotherapy and radiation for breast cancer. Proc Natl Acad Sci U S A 105, 18490–18495 (2008).

35. Boelens, M. C. et al.. Exosome transfer from stromal to breast cancer cells regulates therapy resistance pathways. Cell 159, 499–513 (2014).

36. Fu, N. Y. et al.. Identification of quiescent and spatially restricted mammary stem cells that are hormone responsive. Nat Cell Biol 19, 164–176 (2017).

37. Wellberg, E. A. et al.. Modulation of tumor fatty acids, through overexpression or loss of thyroid hormone responsive protein spot 14 is associated with altered growth and metastasis. Breast Cancer Research 16, (2014).

38. Wu, X. et al.. FASN regulates cellular response to genotoxic treatments by increasing PARP-1 expression and DNA repair activity via NF-κB and SP1. Proc Natl Acad Sci U S A 113, E6965–E6973 (2016).

39. Dekkers, J. F. et al.. High-resolution 3D imaging of fixed and cleared organoids. Nat Protoc 14, 1756–1771 (2019).

40. Rosenbluth, J. M. et al.. Organoid cultures from normal and cancer-prone human breast tissues preserve complex epithelial lineages. Nat Commun 11, (2020).

41. Wolf, F. A., Angerer, P. & Theis, F. J. SCANPY: large-scale single-cell gene expression data analysis. Genome Biol 19, (2018).

42. Traag, V. A., Waltman, L. & van Eck, N. J. From Louvain to Leiden: guaranteeing well-connected communities. Sci Rep 9, (2019).

