## Supplementary figures and images for "A transcriptional response to replication stress selectively expands a subset of *BRCA2*-mutant mammary epithelial cells"

### Supplemental Figure 1

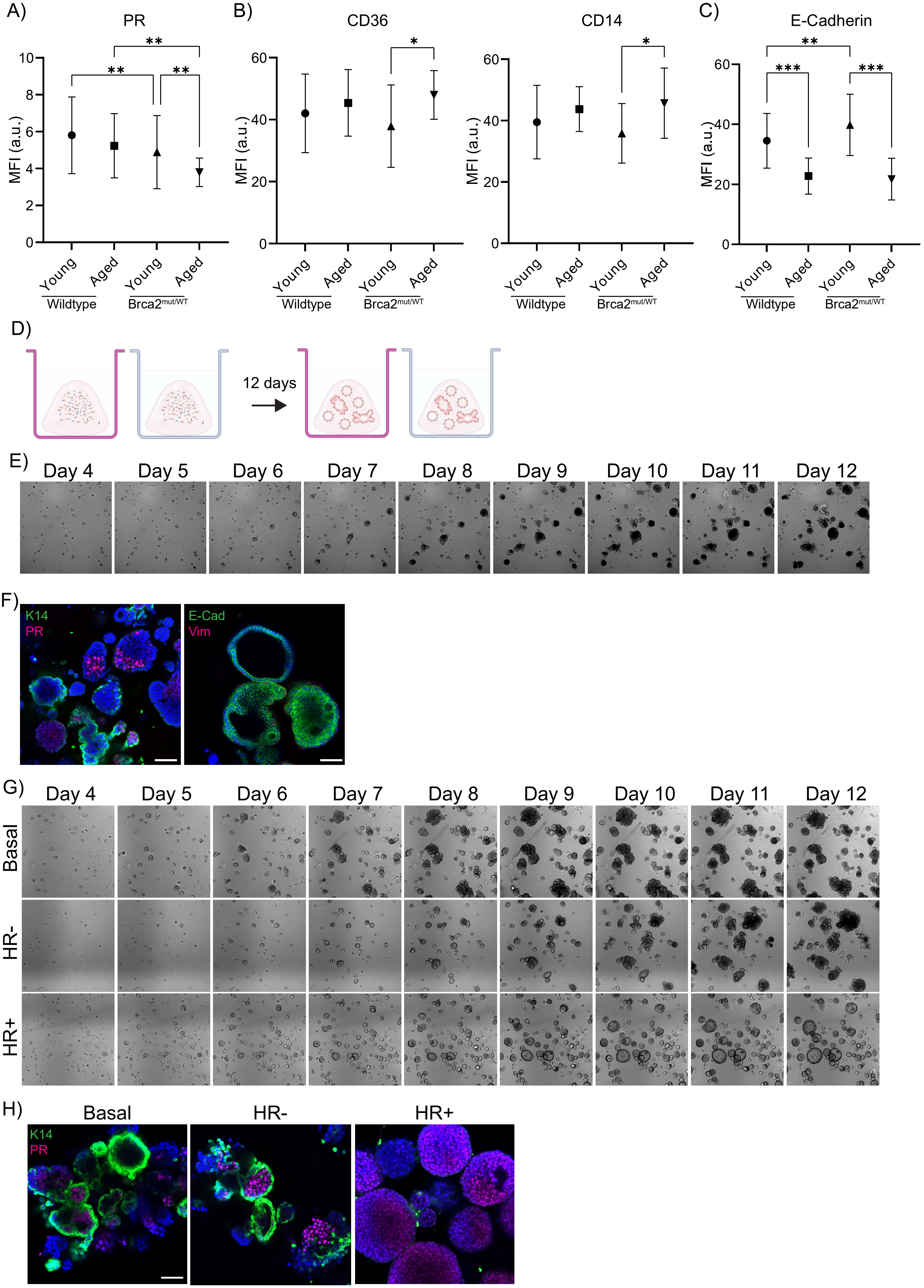

### Supplemental Figure 2

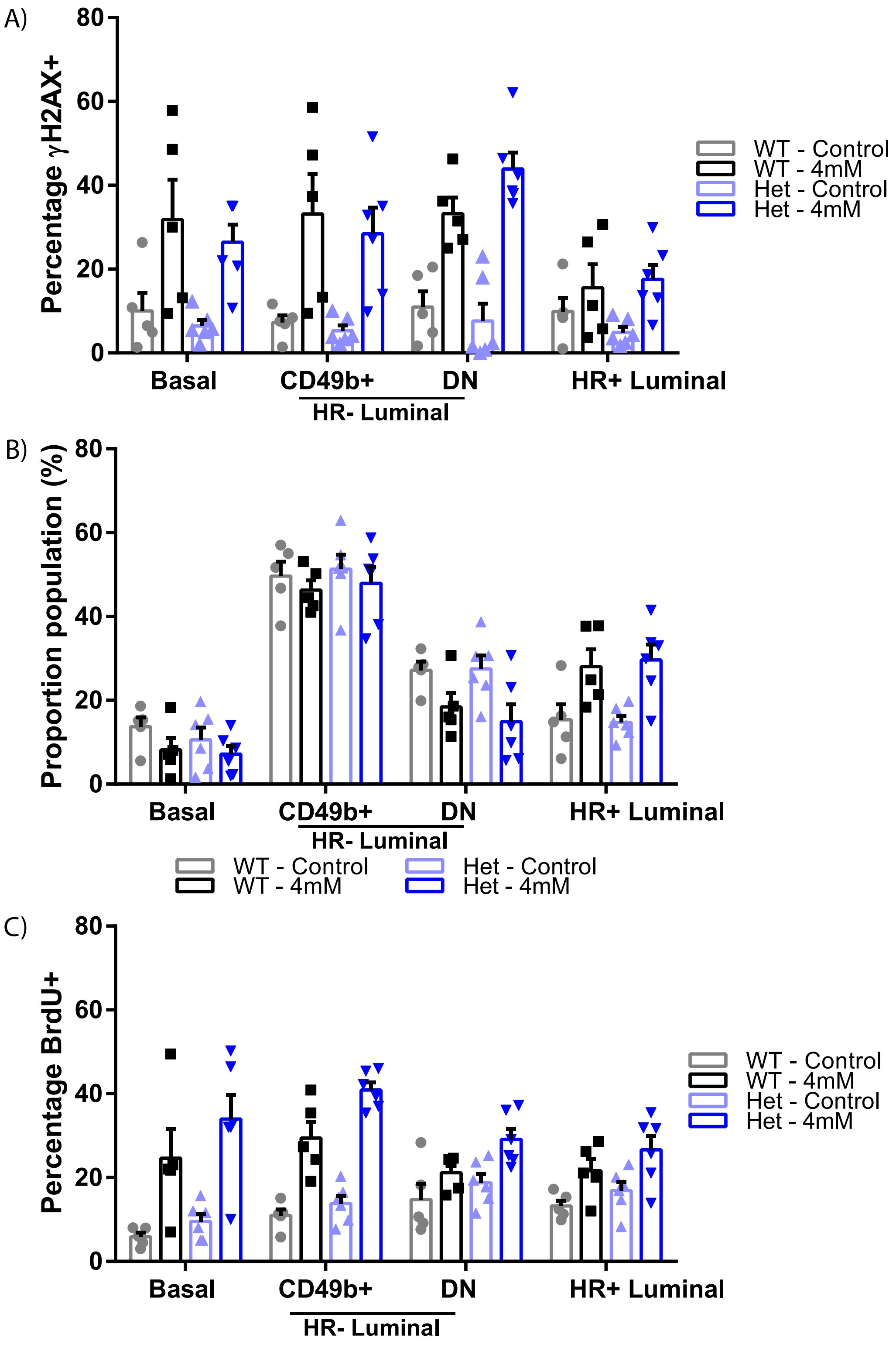


Supp Figure 2.

### Supplemental Figure 3

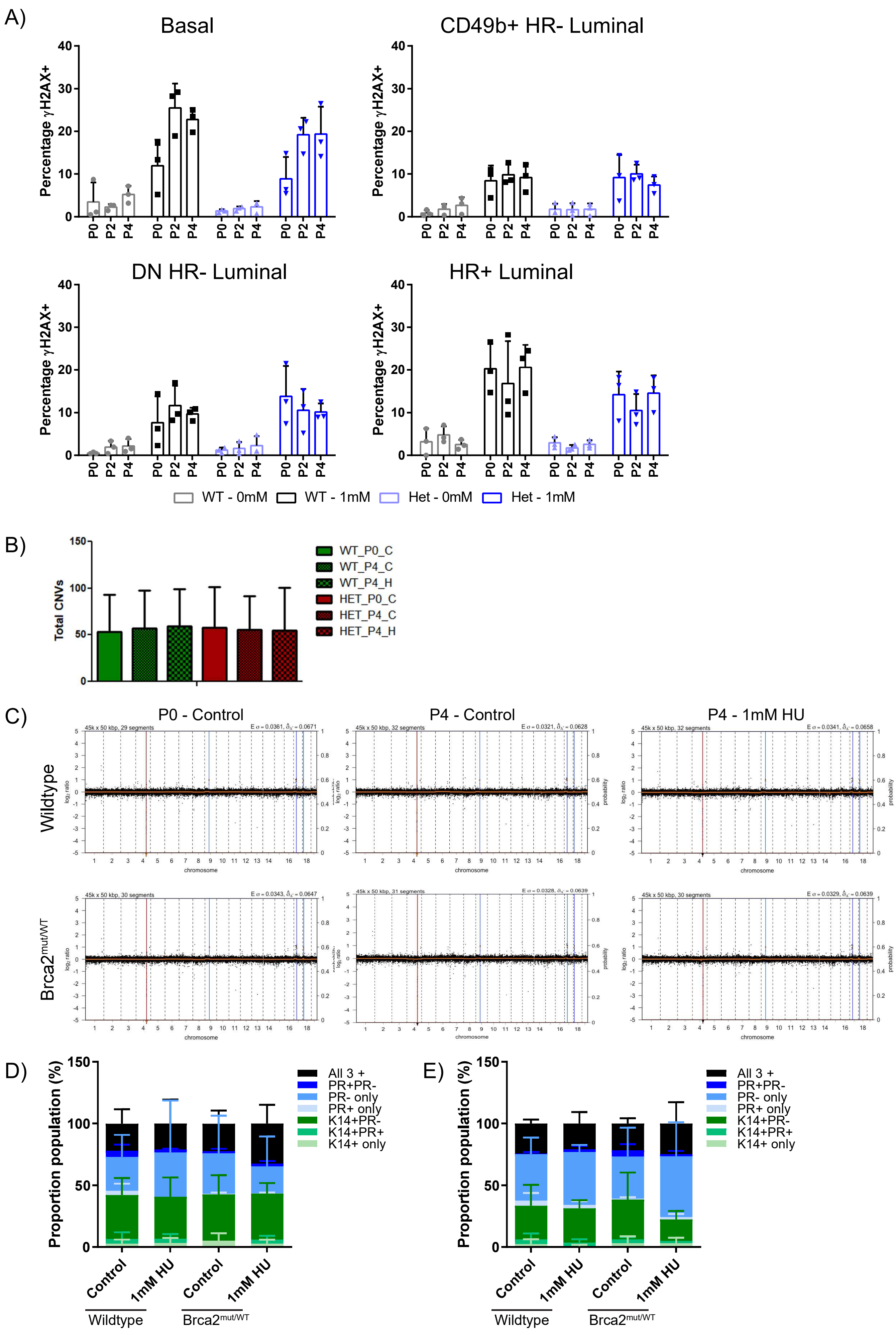


Supp Figure 3.

### Supplemental Figure 4a

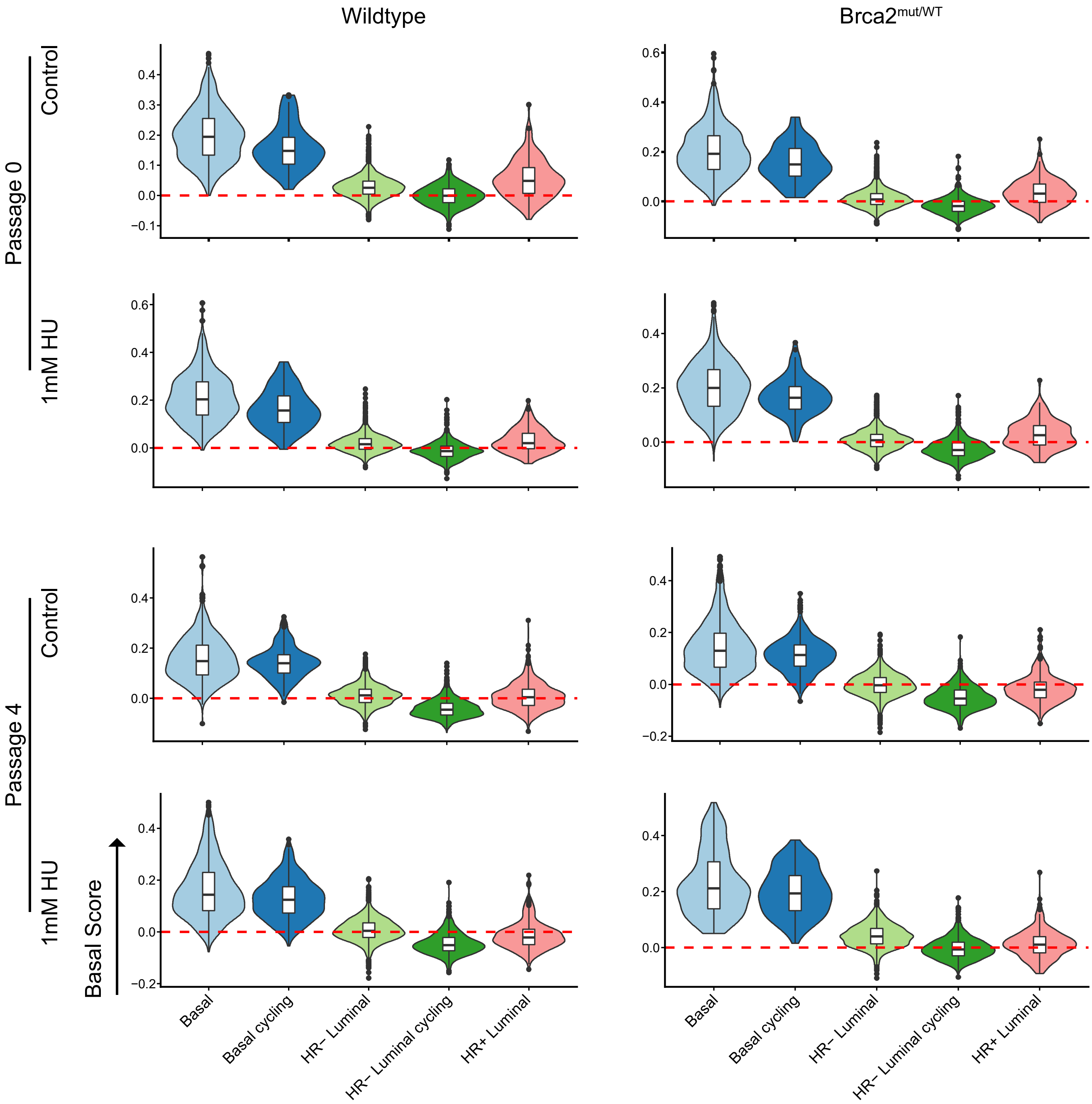

### Supplemental Figure 4b

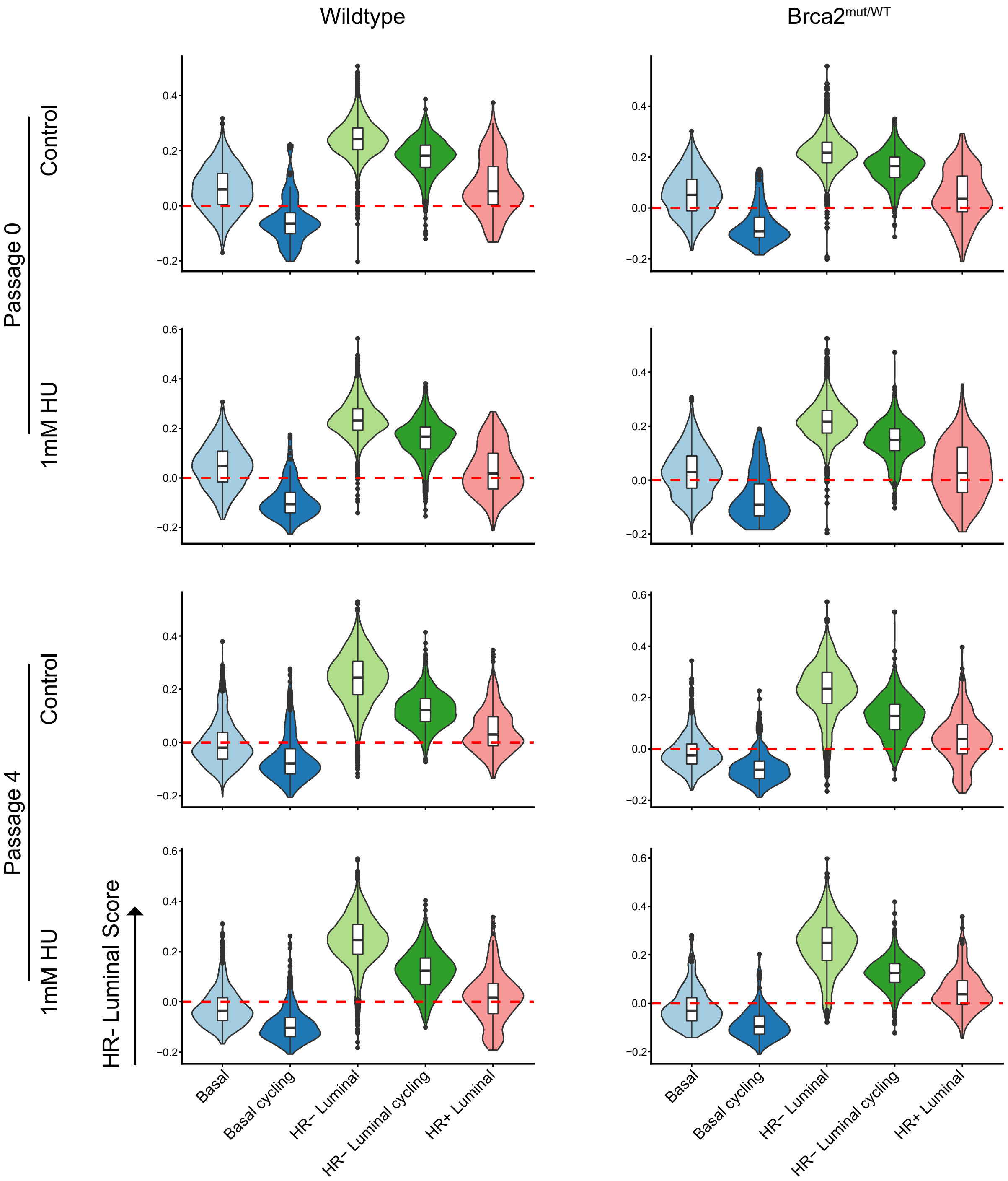

### Supplemental Figure 4c

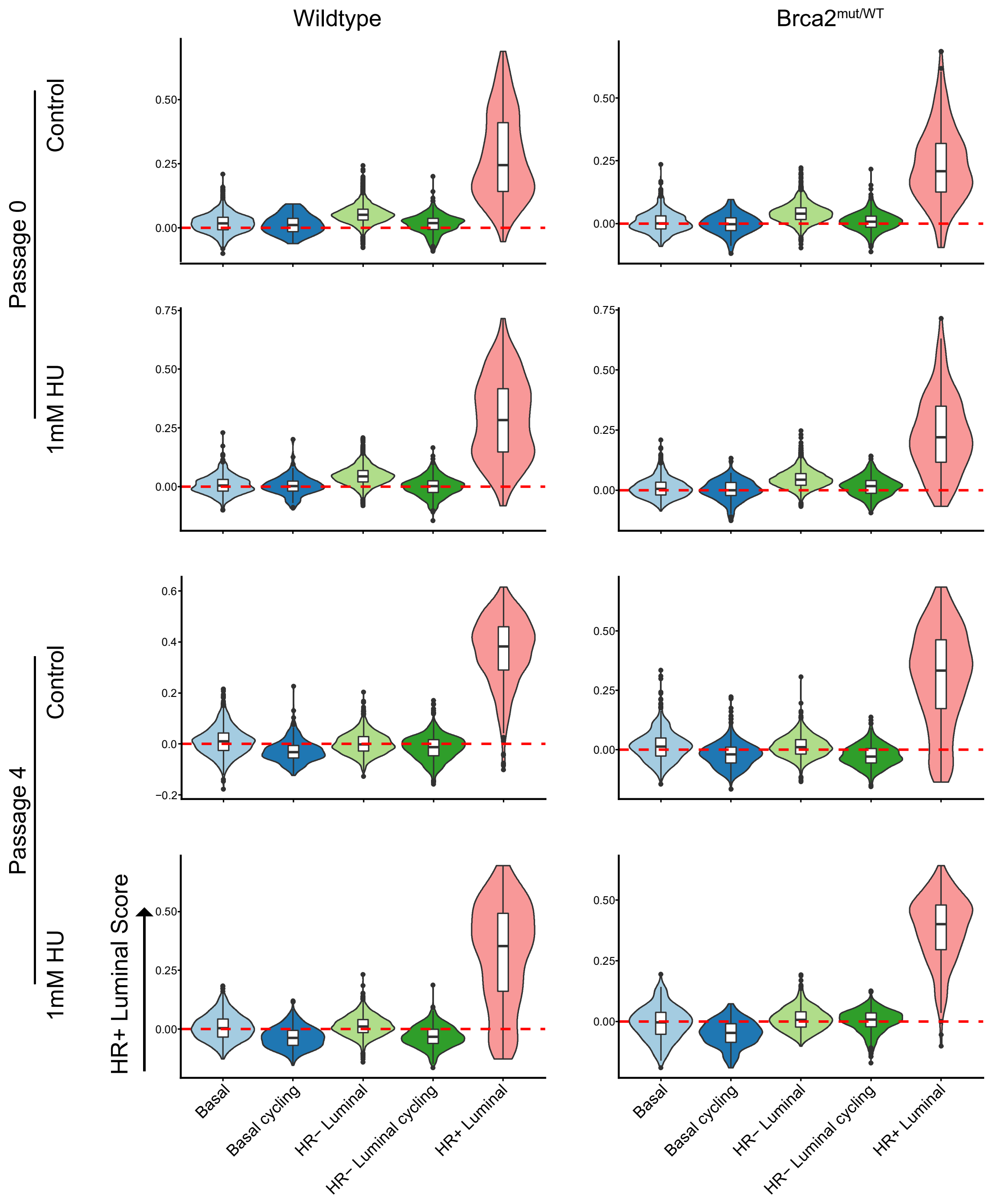

### Supplemental Figure 5

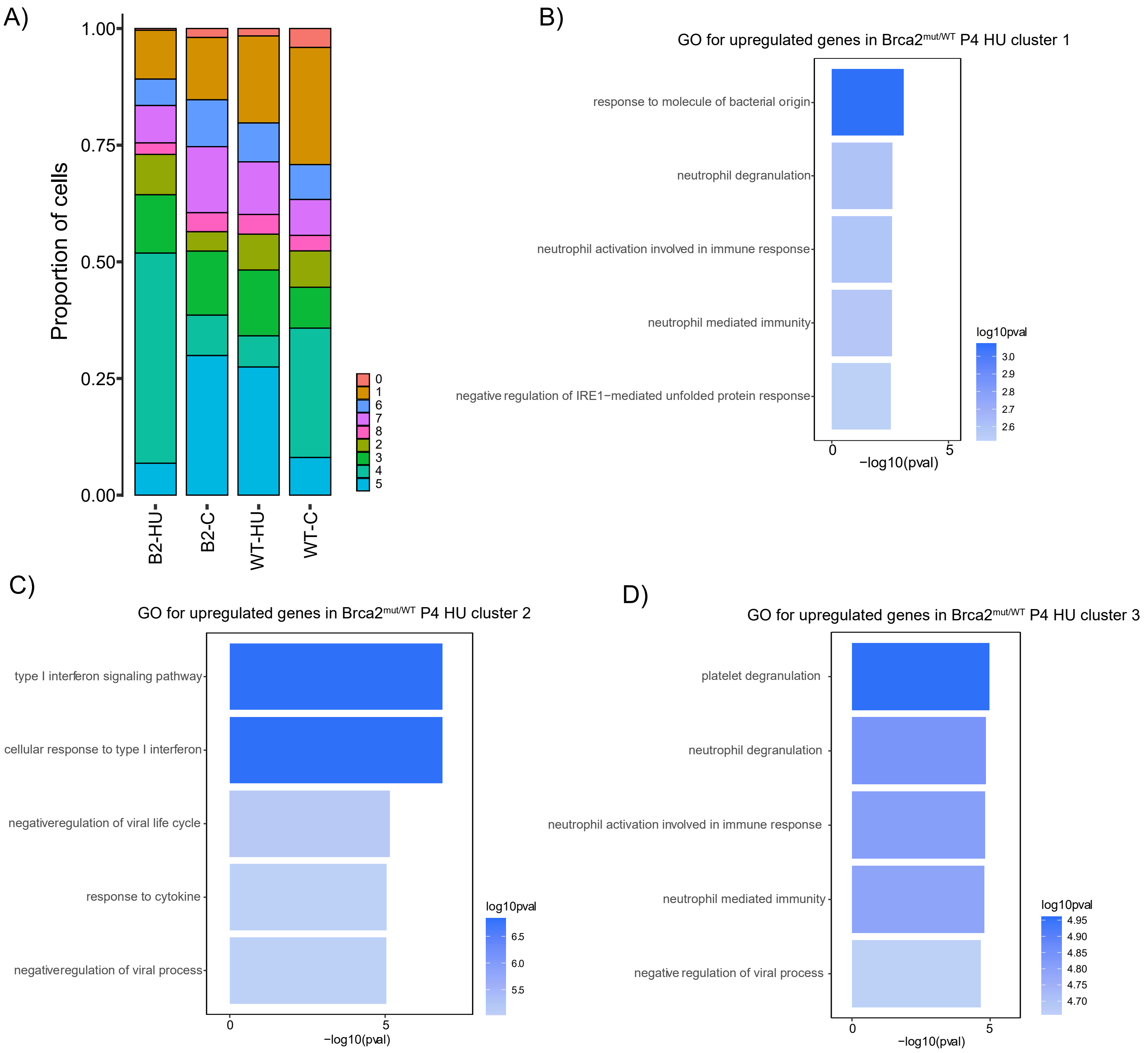

### Supplemental Figure 6

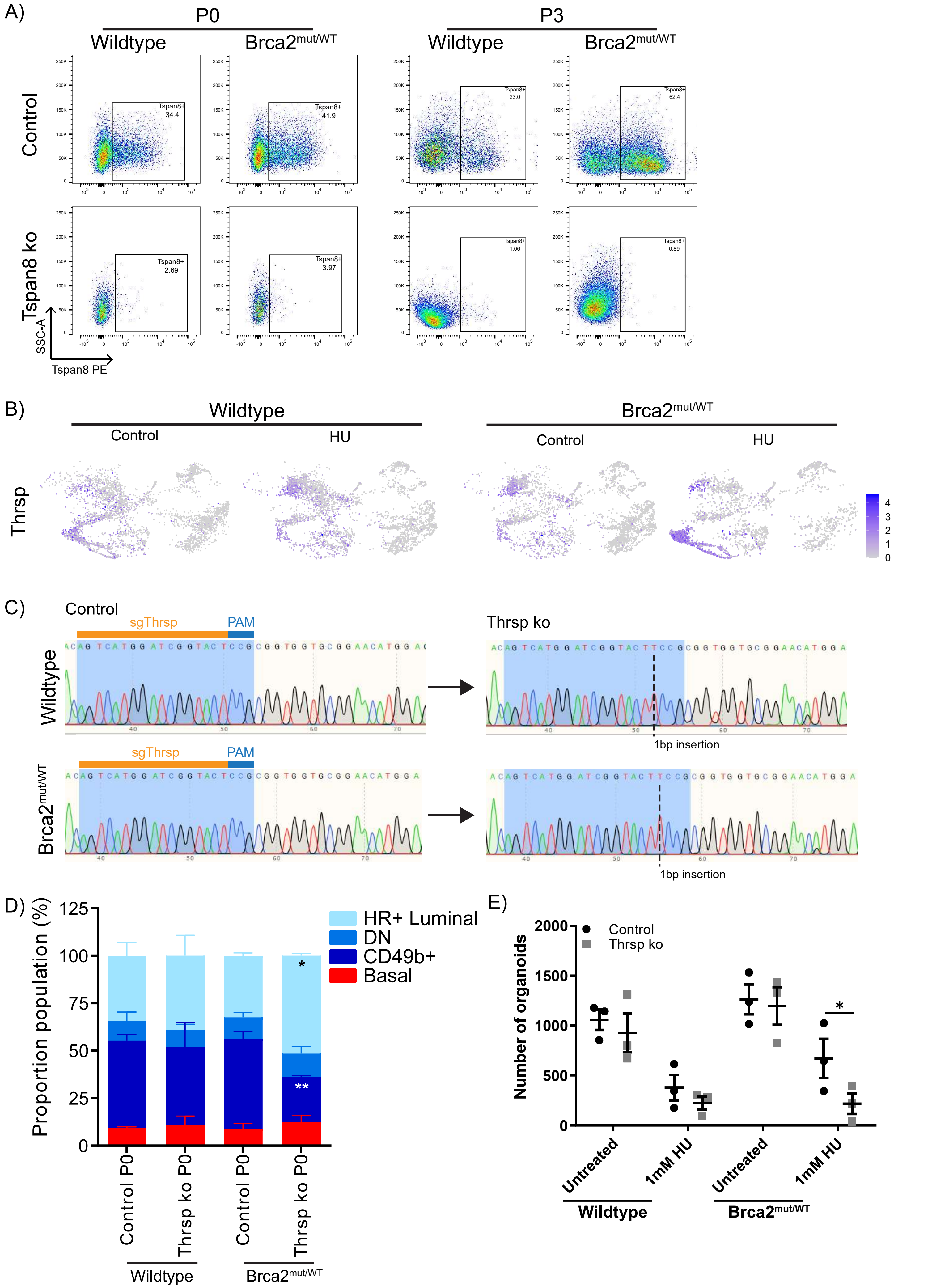
